# The anti-inflammatory potential of helminth-derived peptides/polypeptides: A systematic review in cellular models of inflammation

**DOI:** 10.1101/2025.08.01.668098

**Authors:** Sienna Stucke, Aonghus Feeney, Richard Lalor, Sheila Donnelly, John Pius Dalton, Declan McKernan, Eilís Dowd

**Author notes:** **Corresponding author:** Professor Eilís Dowd.

## Abstract

Helminths are parasitic worms that secrete a plethora of immune regulatory molecules which allow them to dampen inflammatory responses by their host’s immune system to ensure their survival within the host. Their ability to have a compatible existence with their host has led to research into the potential therapeutic effects of helminth-derived molecules for suppression of overactive immune and inflammatory responses in a wide variety of diseases. This systematic review aims to synthesize the published data on helminth-derived peptides/polypeptides (HDPs) with a focus on determining the extent to which they modulate the inflammatory response in *in vitro* cellular models of inflammation. In accordance with PRISMA 2020 guidelines, a predefined systematic search of the PubMed, Web of Science and Medline databases identified relevant studies published up to August 2025, and 79 articles were included after screening. We found that most published studies used LPS or Concanavalin A stimulated macrophages, peripheral blood mononuclear cells or dendritic cells as the cellular model of inflammation. Twenty helminth species from which >60 isolated HDPs were derived were tested in these models, with the nematodes, *Haemonchus contortus* and *Acanthocheilonema viteae*, and the trematode, *Fasciola hepatica,* the most explored species. A common property of these molecules was to ability to significantly reduce the expression or production of pro-inflammatory cytokines such as IL-12, IL-1β, IL-6 and TNF, and significantly increase the expression or production of anti-inflammatory cytokines such as IL-10, TGFβ and IL-4. The effects on other cytokines, including IFNγ which is known to have both pro- and anti-inflammatory effects, were less consistent, with HDPs either decreasing or increasing the levels of this cytokine. This systematic review synthesizes the existing literature in this field and shows that the HDPs secreted by several helminth species have consistently demonstrated effects though modification of cytokine levels and, as such, have therapeutic potential in conditions in which overactive immune and inflammatory responses play a pathogenic role.

## INTRODUCTION

Helminths are parasitic worms, classified into nematodes (roundworms, whipworms, hookworms and flatworms), trematodes (blood-flukes) and cestodes (tapeworms)^1^. These parasites are able to infect a wide variety of species and can cause mild to severe disease in their hosts. Soil-transmitted helminth infections, considered the most important group of neglected tropical diseases by the World Health Organization (WHO), are typically caused by nematode parasites and can lead to intestinal issues, kidney damage, dermatitis, respiratory issues, allergy symptoms, malnutrition, fatigue and other health issues^2^. The parasite eggs are deposited through human and animal faeces where the larvae can survive in water or soil for weeks until they are ingested and then migrate into the intestines, liver, or other tissues where they mature, develop and lay eggs to complete the growth cycle^3^. It is estimated that over 1.5 billion people around the world are infected with one or more of these helminth species, with infections affecting those in developing countries at higher rates due to poor sanitation facilities and infected drinking water^2^.

Interestingly, in developed or high-income countries where worm infections are low, there has been notable increases in immune-mediated conditions such as colitis, allergy and eczema in recent decades that align with patterns of decreasing helminth infection^4,5^. This inverse correlation between helminth infection and immune/inflammatory disease suggests that by limiting exposure to the diverse pathogens through safer hygiene and sanitation practices, more humans today lack the immune-protective effects provided by helminth infection against inflammatory and allergenic diseases^6^. This idea has evolved into the ‘old friends’ hypothesis, proposing that, because helminths evolved alongside the adaptive immune system they provide some protection against certain diseases, and in their chronic infective state, they are more like ‘old friends’ than foe^7^. Indeed, in endemic populations it seems the ubiquitous presence of helminths has even impacted the immune response at the genome level with evidence of SNP in genes associated with the Th2 immune response^8^. These observations have stimulated extensive research into elucidating the molecular mechanisms behind the immunomodulatory properties of worms, and particularly the molecules they secret as a potential novel source of biotherapeutics for the treatment of immune and inflammatory conditions.

It is now known that helminths secrete a plethora of immunosuppressive molecules which allow then to remain undetected for long periods of time by their host^9^. These helminth-derived peptides/polypeptides (HDPs) are immunomodulatory and can modulate the host’s immune response by shifting the balance between the Th1 and Th2 responses. This leads to an immunosuppressive state during the parasite’s chronic infective stage, allowing them to remain tolerated by the host for months, years or even decades. While HDPs are secreted for parasite protection inside the host, because they are immunosuppressive, they may have therapeutic potential as isolated (or synthetic) peptides/polypeptides in a wide variety of diseases in which overactive immune and inflammatory responses play a pathogenic role^1,10^. Multiple types of HDPs from a variety of helminth species have been isolated and purified (or synthesized) in the past twenty years. These have been shown to modulate the immune and inflammatory response in multiple cell types, particularly through altered cytokine responses. However, to date, there has been no systematic review of this literature, and thus, this review aims to systematically synthesize the published data on HDPs and the extent to which they modulate the inflammatory response in cellular models of inflammation.

## METHODS

### Search strategy

This study was completed in accordance with the PRISMA 2020 guidelines^11^ to find articles in which HDPs were assessed in cellular models of inflammation. The literature search was completed in the PubMed, Web of Science and Medline databases with the specific search string: helminth AND secret* AND (immunomod* OR immunosup* OR antiinflam*). This search identified a total of 907 records, spanning July 2001 to August 2025. These articles were then screened according to the strategy outlined below and depicted in the PRISMA flow diagram (Fig. 1). This yielded 79 articles which were included in this systematic review. All records were managed in the Endnote and Microsoft Excel software packages.

### Screening strategy

Once all duplicate records were removed, the remaining articles were screened by title and abstract according to the following inclusion and exclusion criteria. Inclusion criteria: (1) original research study, (2) HDPs, (3) cellular inflammatory model. Exclusion criteria: (1) review article, (2) not helminth-derived (3), undefined helminth-derived molecule, (4) not peptide helminth-derived molecule, (5) not tested in inflammatory model and (6) not peer-reviewed or redacted. In the full-text screen, three other exclusion criteria were applied (7) no cytokine measure, (8) no appropriate control and (9) not in a cellular (*in vitro*) model. After screening, 63 articles met the inclusion and exclusion criteria, and a further 16 articles were added from the cited publications in the selected articles. Thus, a total of 79 articles were included in this systematic review (Table 1 and Supplementary Excel file).

**Fig. 1.**
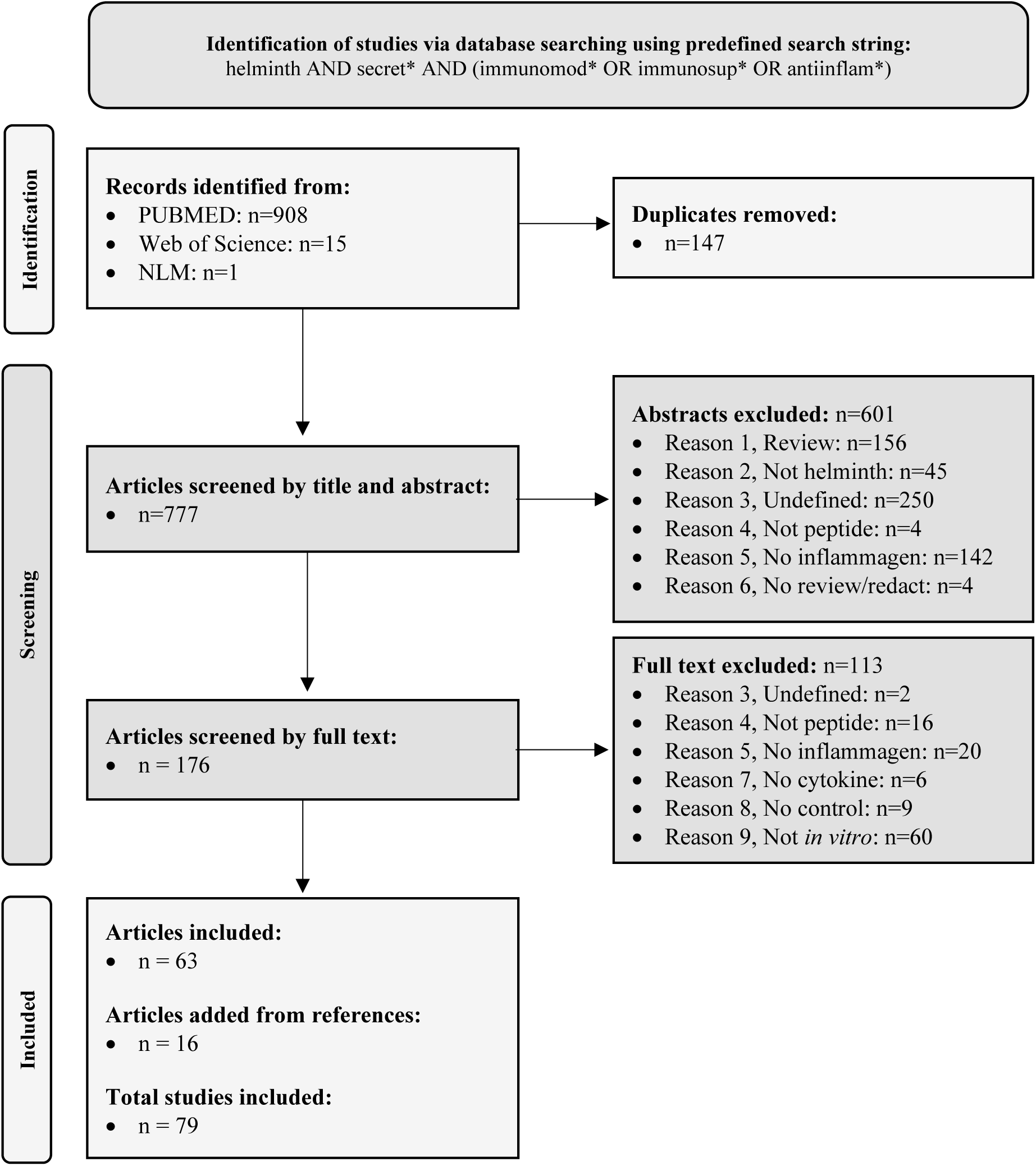
PRISMA flow chart. Flow chart based on PRISMA 2020 guidelines^11^ detailing the screening strategy employed for the study selection in the present study.

### Data extraction

Variables manually extracted from the selected articles included cell type, inflammagen, helminth species and HDP name, as well as cytokines measured (Table 1 and Supplementary Excel file). Because any given article may have had several experimental parameters (e.g. different HDP concentrations or different incubation times etc.), to comprehensively synthesize this literature, each of these was considered a separate “record” for cytokine data analysis. Cytokine levels were measured by qPCR, Western immunoblotting or ELISA, and expressed as relative values or concentration as appropriate. Specific values were extrapolated from the relevant figure(s) within each article using GetData Graph Digitizer software. The effects of the HDPs were then assessed by calculating the percent change in cytokine levels when cells were challenged with inflammagen in the absence versus the presence of the HDPs. Data extraction was completed independently by two of the authors (SS and AF) and cross-checked for accuracy.

### Statistical analyses

General study characteristics are shown as pie charts. All data related to percent cytokine change are shown as scatter plots depicting individual records with the mean ± standard error of the mean (SEM). To determine if the HDPs reduced or increased cytokine levels beyond zero, data were analysed using one-sample t-test (with the hypothesised population mean set to zero). To determine the effect of cell type and inflammagen on the efficacy of HDPs, the data were analysed by two-way ANOVA with *post-hoc* Bonferroni.

## RESULTS

### General study characteristics

Seventy-nine articles were included in this review in which HDPs were assessed for their immunomodulatory effects in *in vitro* cellular models of inflammation (Table 1 and Supplementary Excel file). We found that most published studies used macrophages, peripheral blood mononuclear cells (PBMCs) or dendritic cells as the target immune cell (Fig. 2A), and LPS or Concanavalin A to elicit inflammatory responses (Fig. 2B). Although 20 different helminth species tested in these studies, the most widely used were the nematodes, *Haemonchus contortus* and *Acanthocheilonema viteae,* the trematodes, *Fasciola hepatica*, *Schistosoma mansoni* and *Schistosoma japonicum*, and the cestode, *Echinococcus granuloses* (together tested in 65% of studies) (Fig. 2C). From the collective helminth species, >60 different sequenced HDPs were tested (Table 1 and Supplementary Excel file), and although there was no widely used HDP, ES-62 (a phosphorylcholine-containing glycoprotein secreted by *A. viteae)* was used most frequently (n=7 articles).

**Fig. 2.**
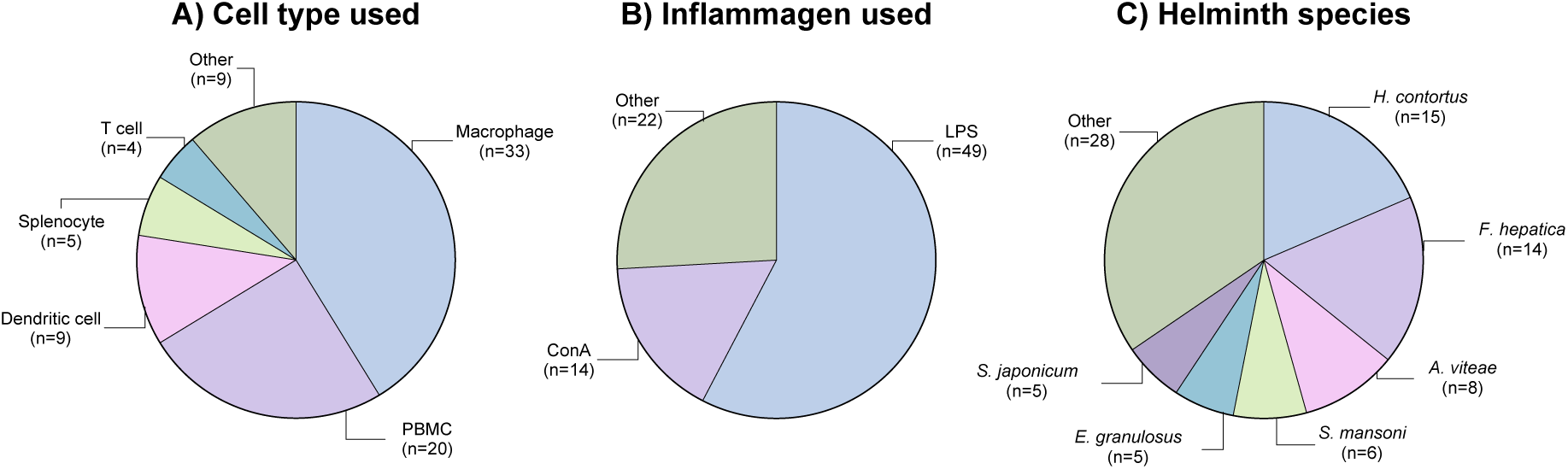
General study characteristics. A total of 79 articles were included in this review and a summary of the key characteristics of these articles is depicted in these pie charts. These show the proportions of these studies using different A) cell types, B) inflammagens and C) helminth species. LPS: lipopolysaccharide; ConA: Concanavalin A; PBMC: peripheral blood mononuclear cell.

**Table 1.**
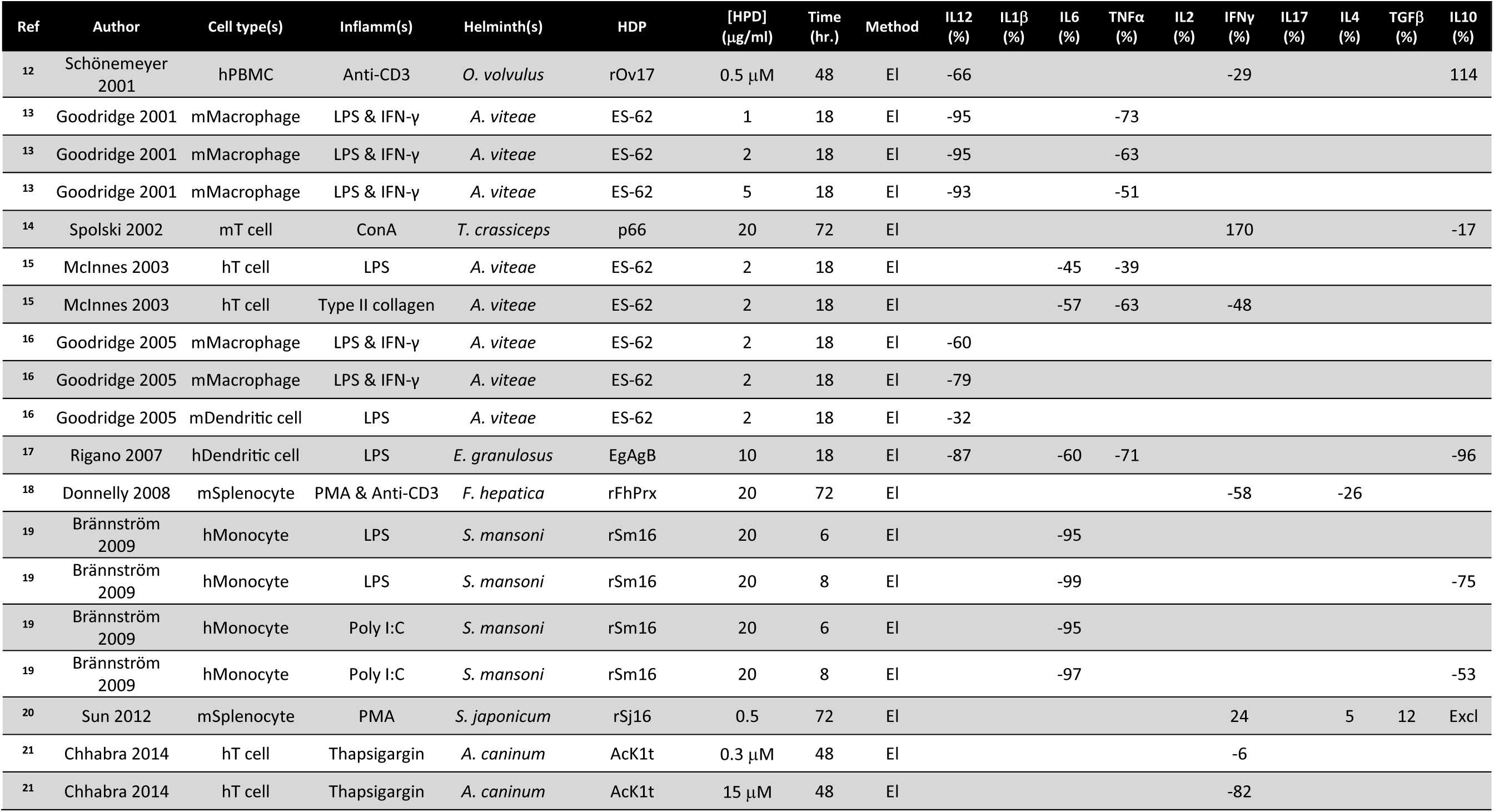
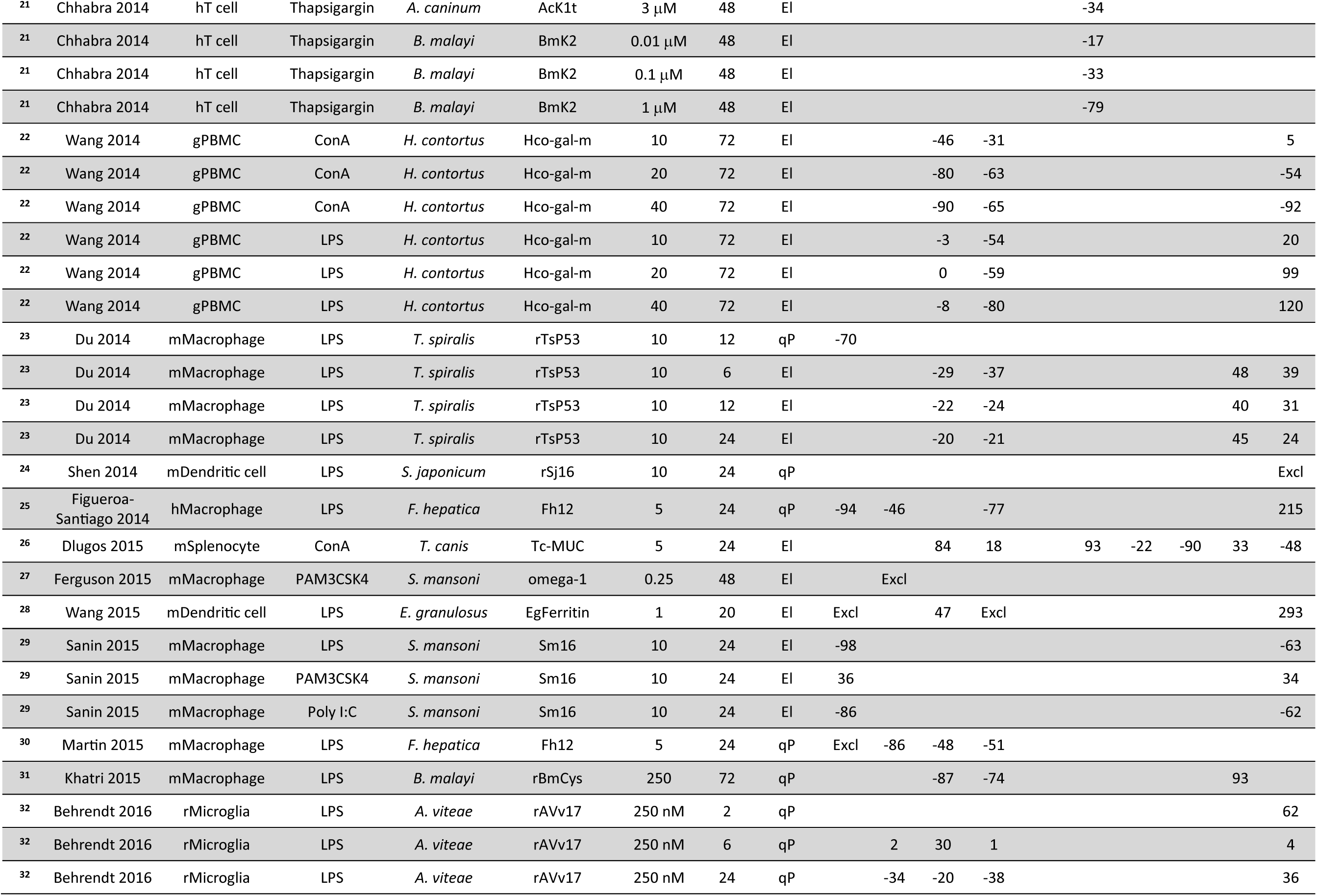
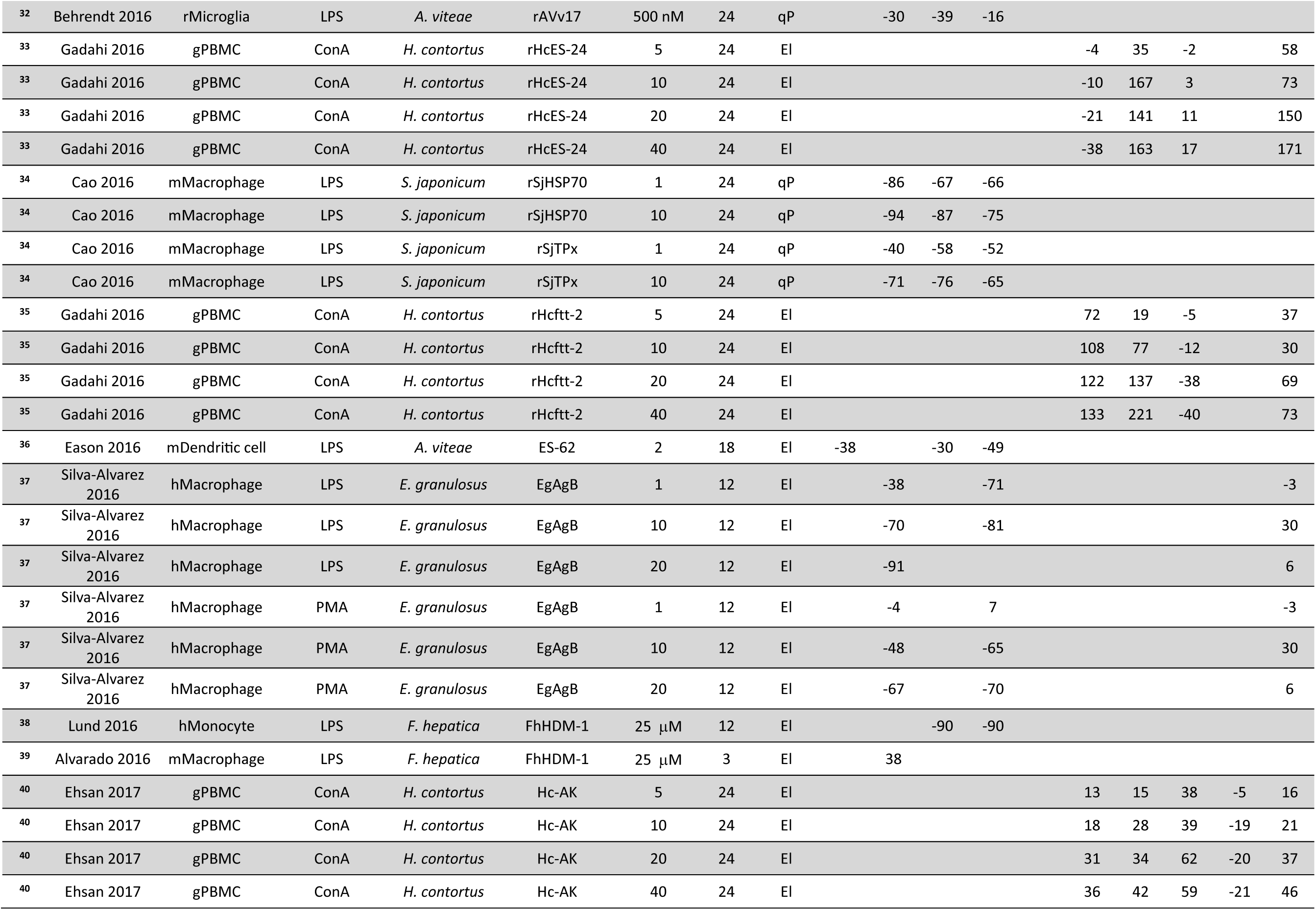
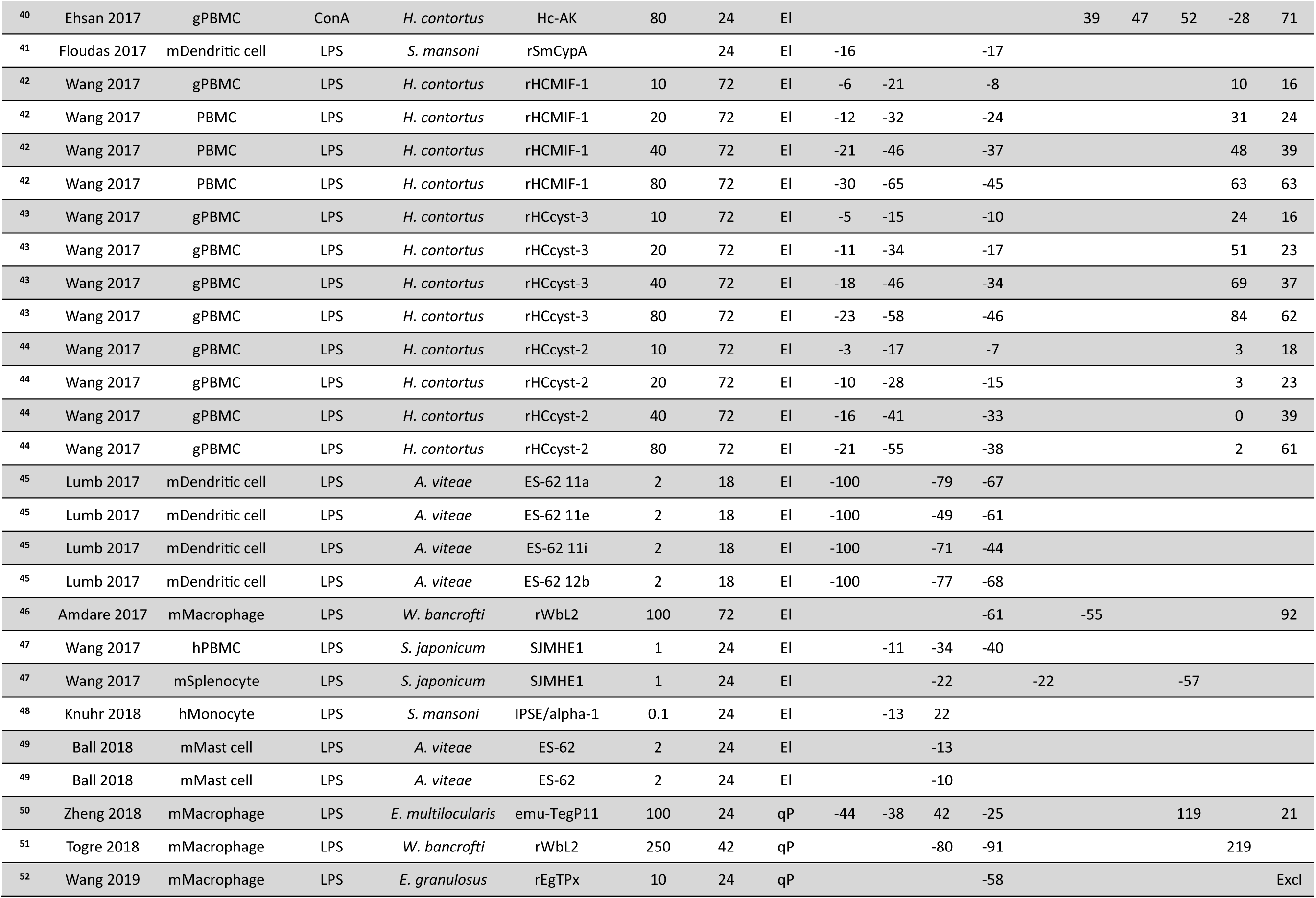
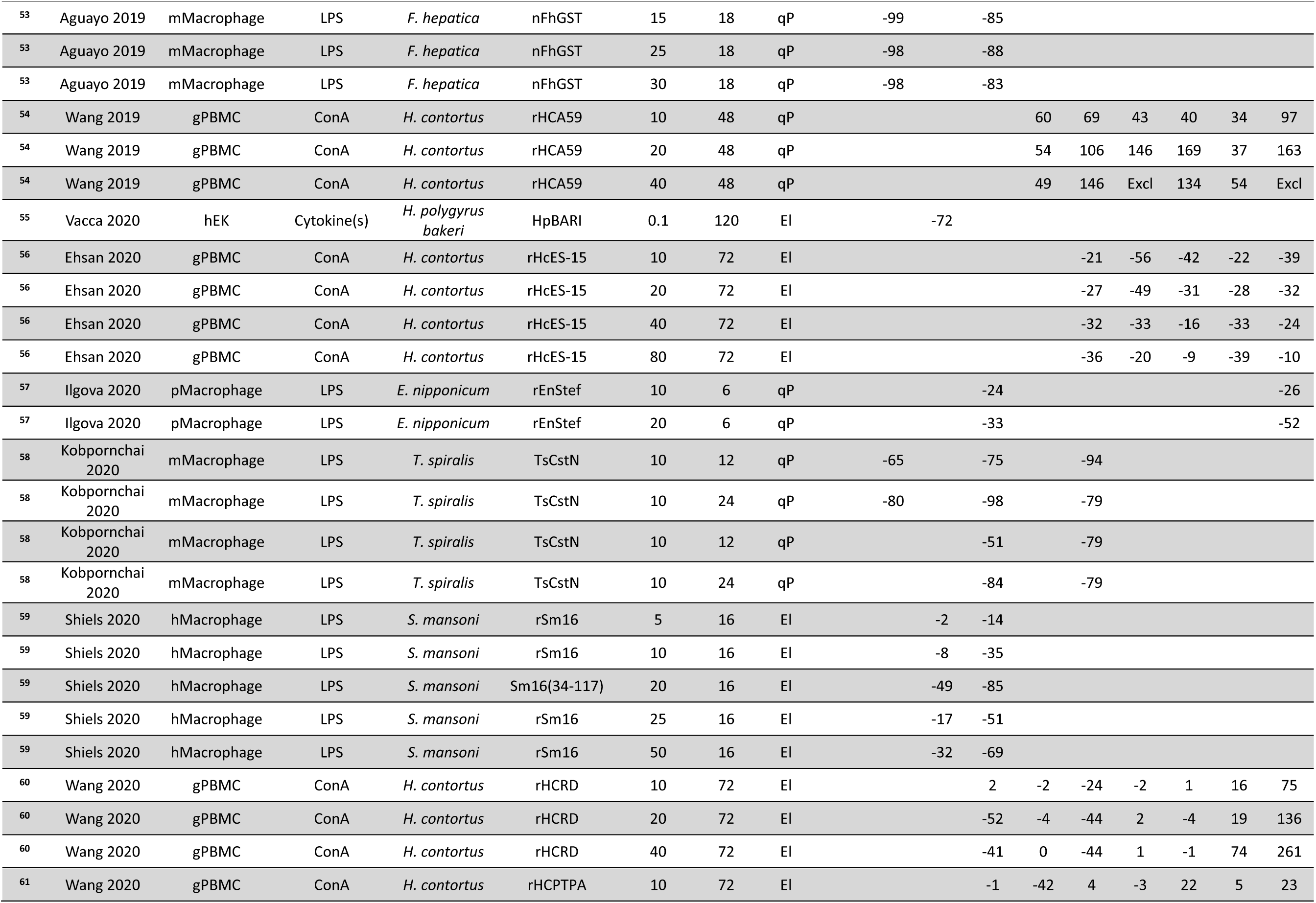
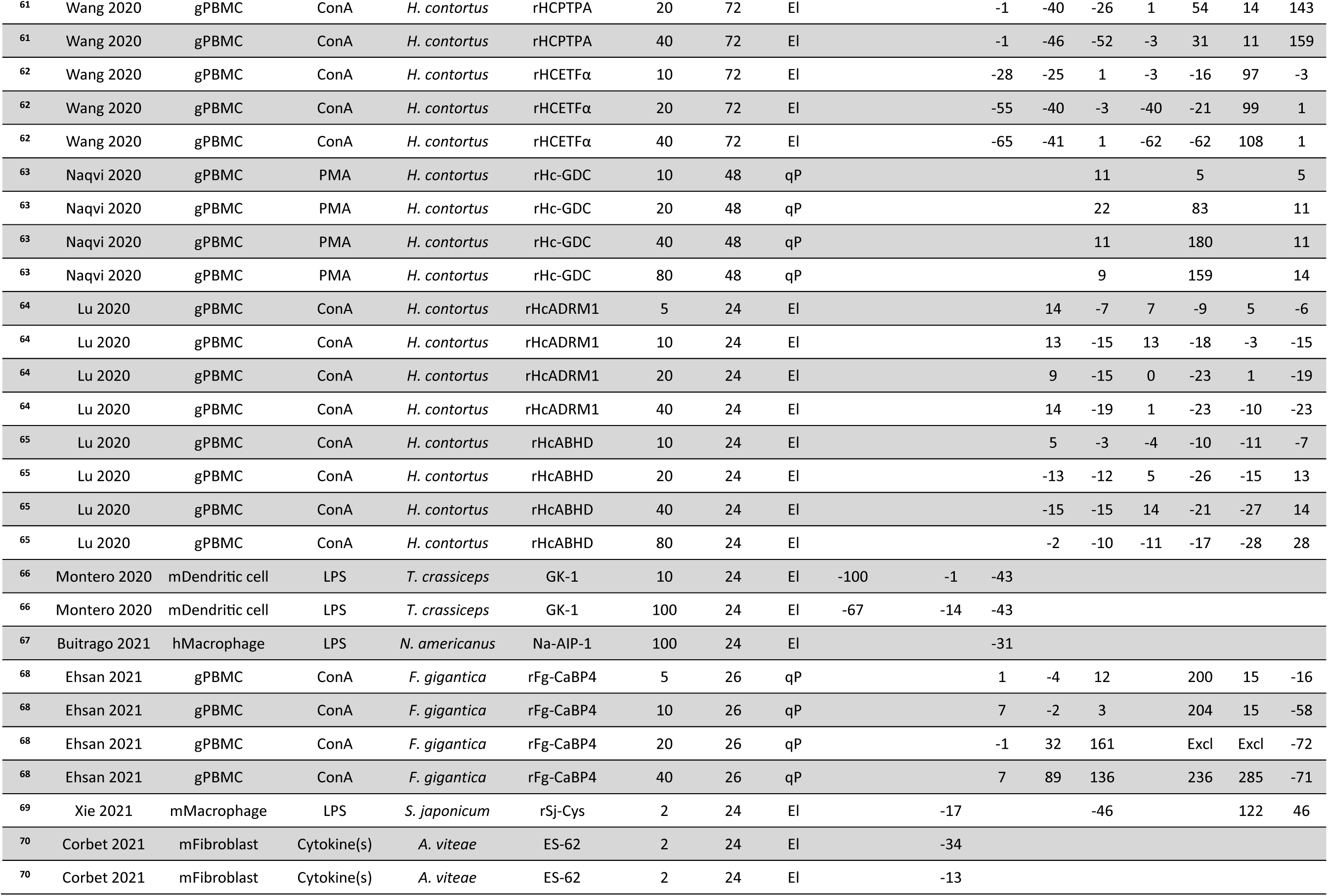
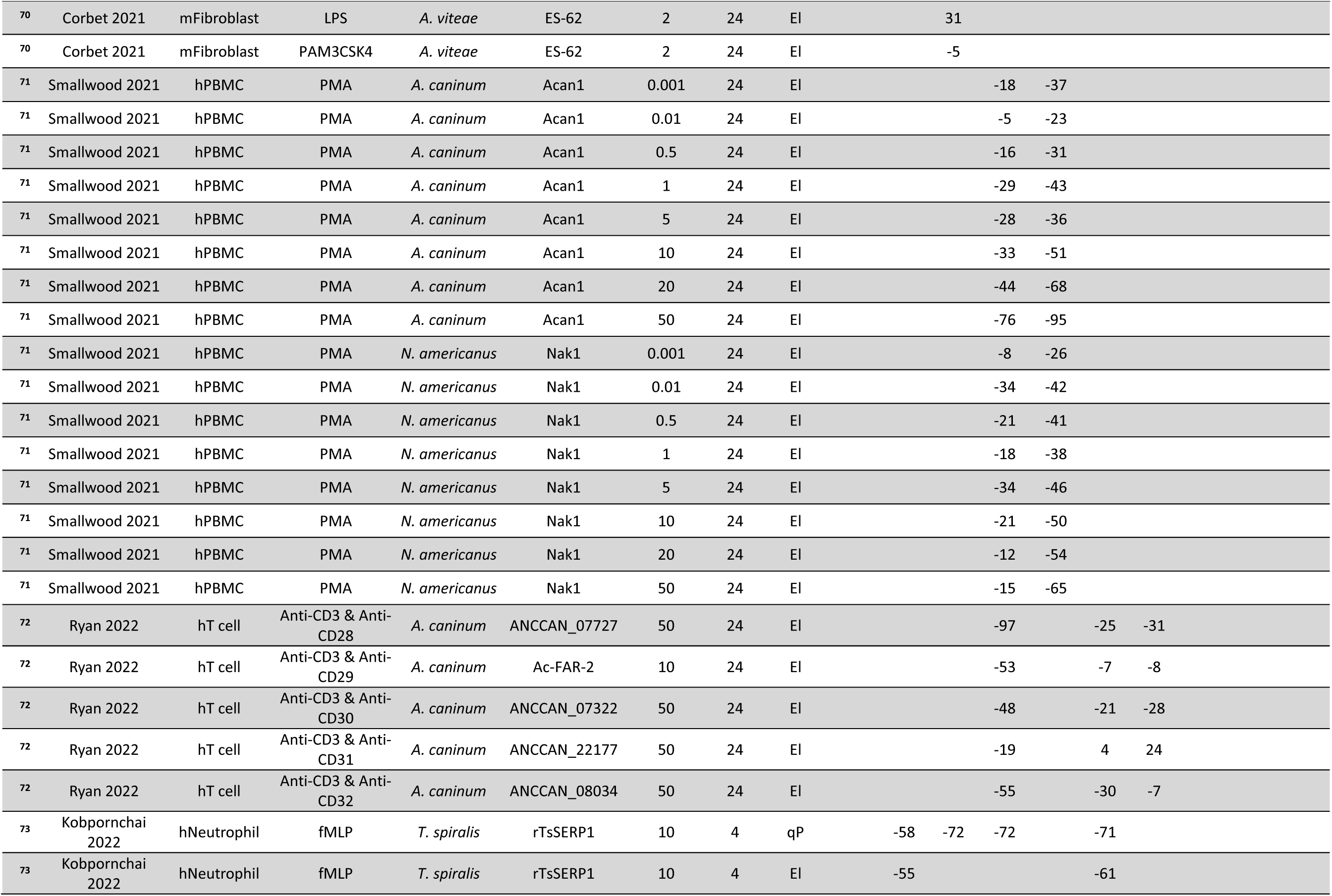
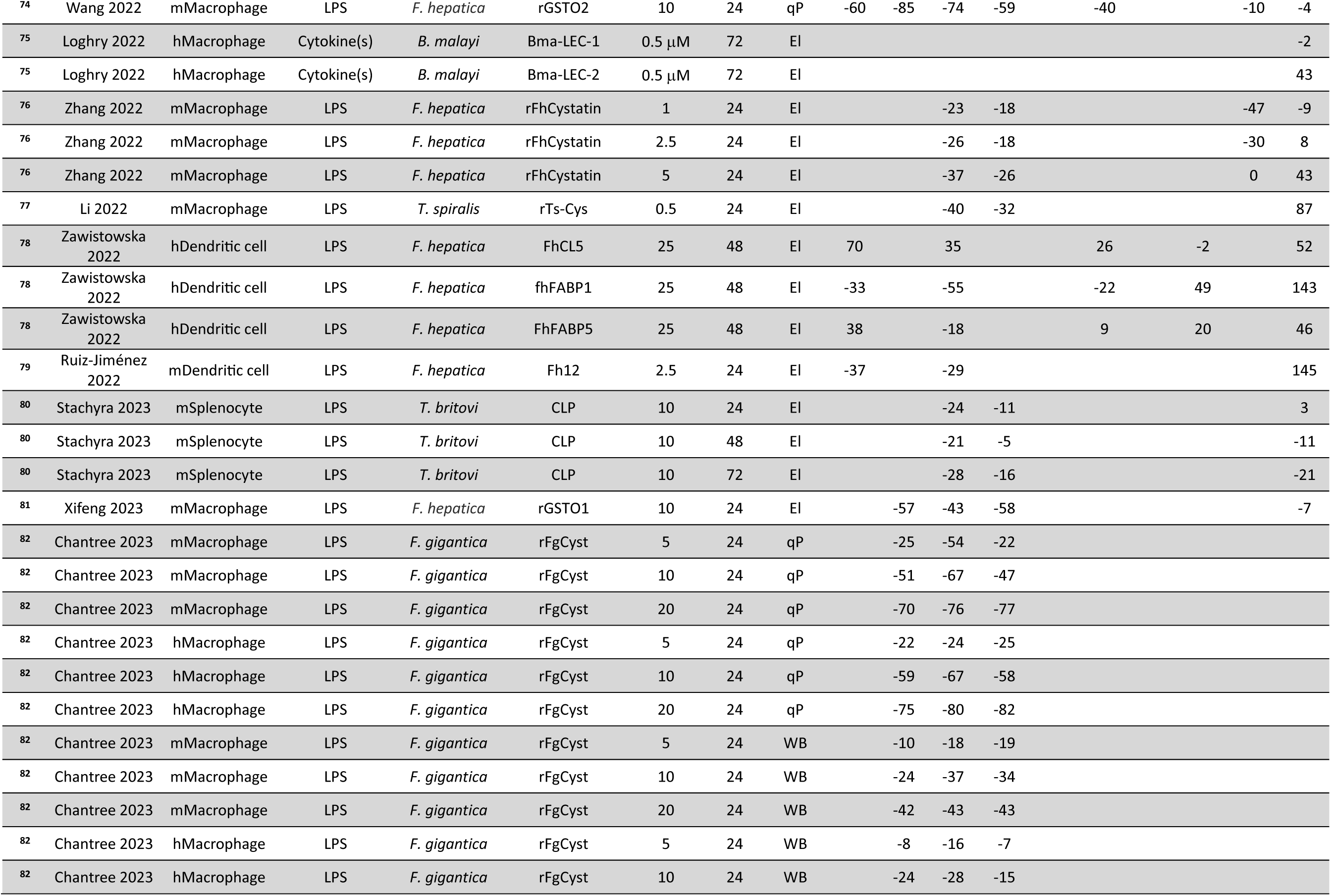
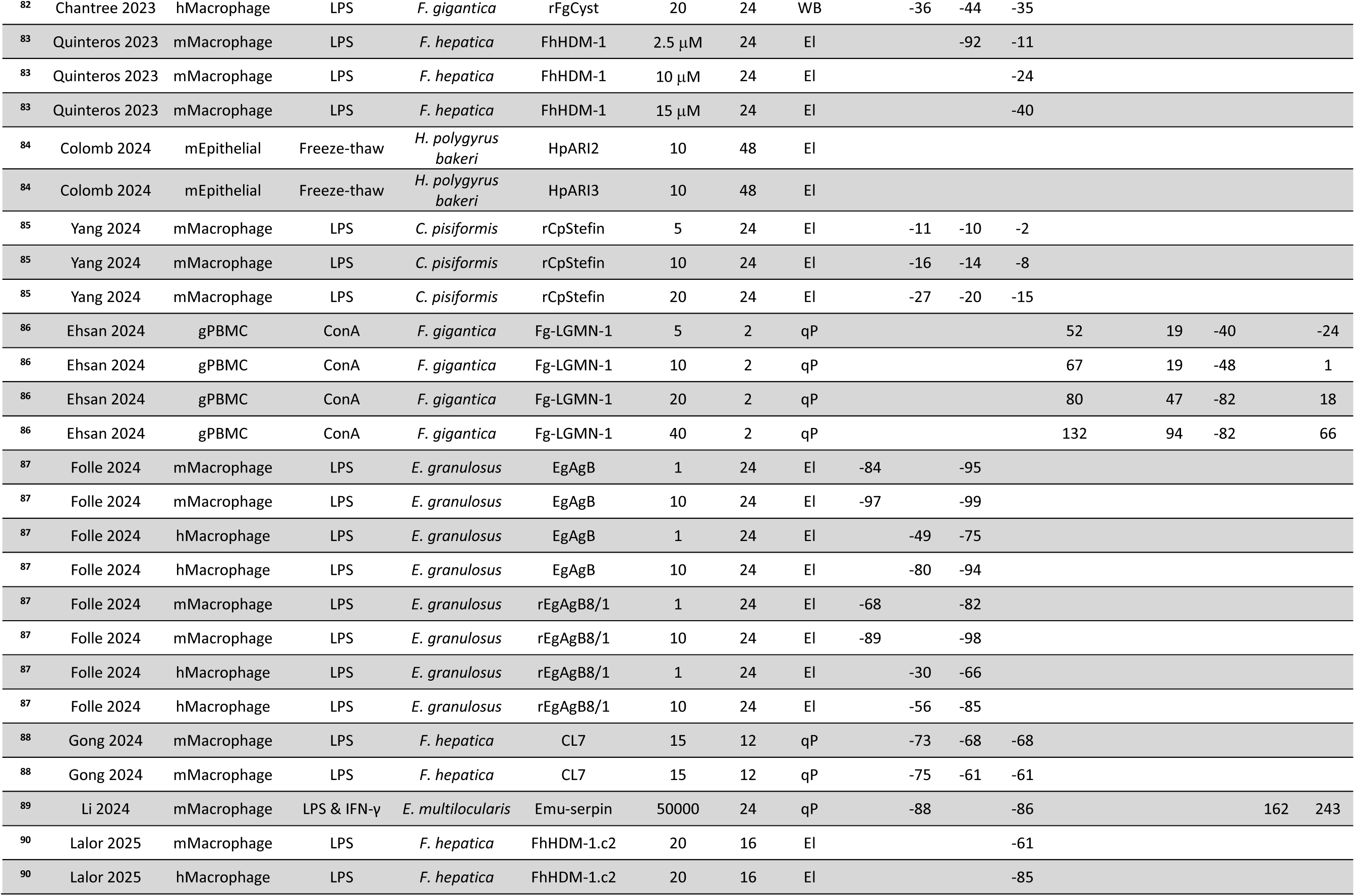
Studies and individual records included in this review. A total of 79 articles were included in this review from which 229 separate records were identified. For each of these records, the percent change in cytokine levels is shown. The lowercase letter before each cell types denotes the species of origin: human (h); mouse (m), goat (g), rat (r), pig (p). Method refers to the method used for cytokine analysis: ELISA (El); qPCR (qP); WB (Western immunoblotting). See Supplementary Excel file for more details and specific values. Abbreviations: ConA: Concanavalin A; EK: embryonic kidney; fMLP: N-Formylmethionine-leucyl-phenylalanine; Inflamm(s): Inflammagen(s); LPS: lipopolysaccharide; PBMC: peripheral blood mononuclear cell; PMA: phorbol myristate acetate. Excl: Values excluded as statistical outliers.

### Effect of HDPs on cytokines

From the 79 articles included in this review, 229 separate records were identified in which HDPs were assessed for their effect on cytokine levels in cellular models of inflammation. Taking all this data together, the effect of the HDPs on the most widely assessed cytokines was first examined (Fig. 3). This revealed that levels of the pro-inflammatory cytokines, IL-12, IL-1β, IL-6 and TNFα were all significantly reduced by the HDPs (IL-12: *t*_(42)_=7.41, *P*<0.0001; IL-1β: *t*_(61)_=12.73, *P*<0.0001; IL-6: *t*_(92)_=10.55, *P*<0.0001; TNFα: *t*_(138)_=17.66, *P*<0.0001). In contrast, levels of the anti-inflammatory cytokines, IL-10, TGFβ and IL-4 were significantly increased (IL-10: (*t*_(109)_=4.80, *P*<0.0001; TGFβ: *t*_(56)_=3.54, *P*<0.001; IL4: *t*_(55)_=2.01, *P*<0.05). The HDPs also increased levels of the pro-inflammatory cytokine IL-17 (t_(45)_=2.78, *P*<0.01), but neither IL-2 nor INFγ were significantly changed (from zero) overall.

**Fig. 3.**
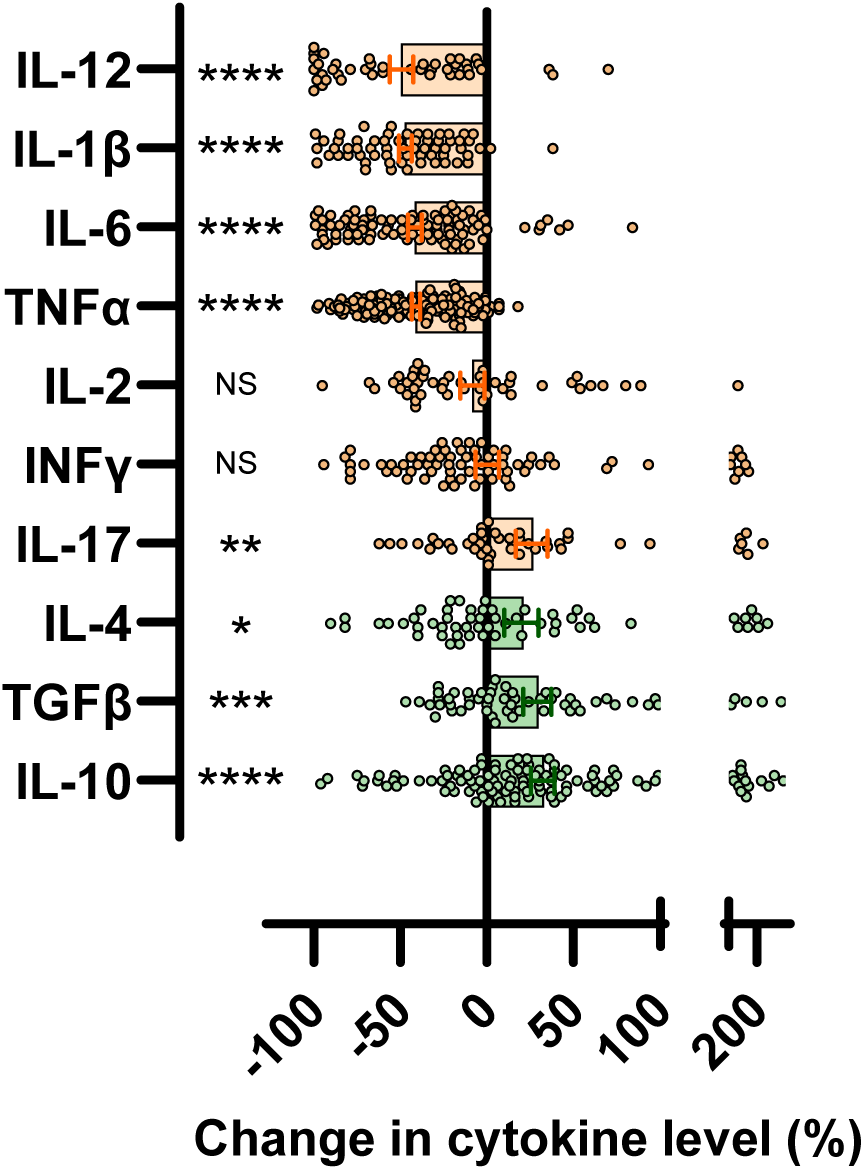
Effect of HDPs on cytokines overall. The effect of the HDPs on the most widely assessed cytokines. Each data point represents a specific record extracted from Table 1/Supplementary Excel file, and the mean±sem is also shown. Data were analysed by one-sample test with the hypothesised population mean set to zero. \*\*\*\**P*<0.0001, \*\*\**P*<0.001, \*\**P*<0.01 and \**P*<0.05. NS: not-significantly different to zero.

### Effect of cell type and inflammagen on efficacy of HDPs

The effects of main cell type and inflammagen on the ability of the HDPs to modify cytokine levels was then assessed (Fig. 4). This highlighted that cell type (Fig. 4A) but not inflammagen (Fig. 4C) had a significant effect on the ability of the HDPs to reduce pro-inflammatory cytokine expression. Specifically, the HDPs reduced IL-12 and TNFα significantly more in macrophages than PBMCs (Fig. 4A; Cell type: *F*_(1, 247)_ = 32.78; *P*<0.0001). In contrast, neither cell type (Fig. 4B) nor inflammagen (Fig. 4D) affected the ability of the HDPs to increase anti-inflammatory cytokine levels.

**Fig. 4.**
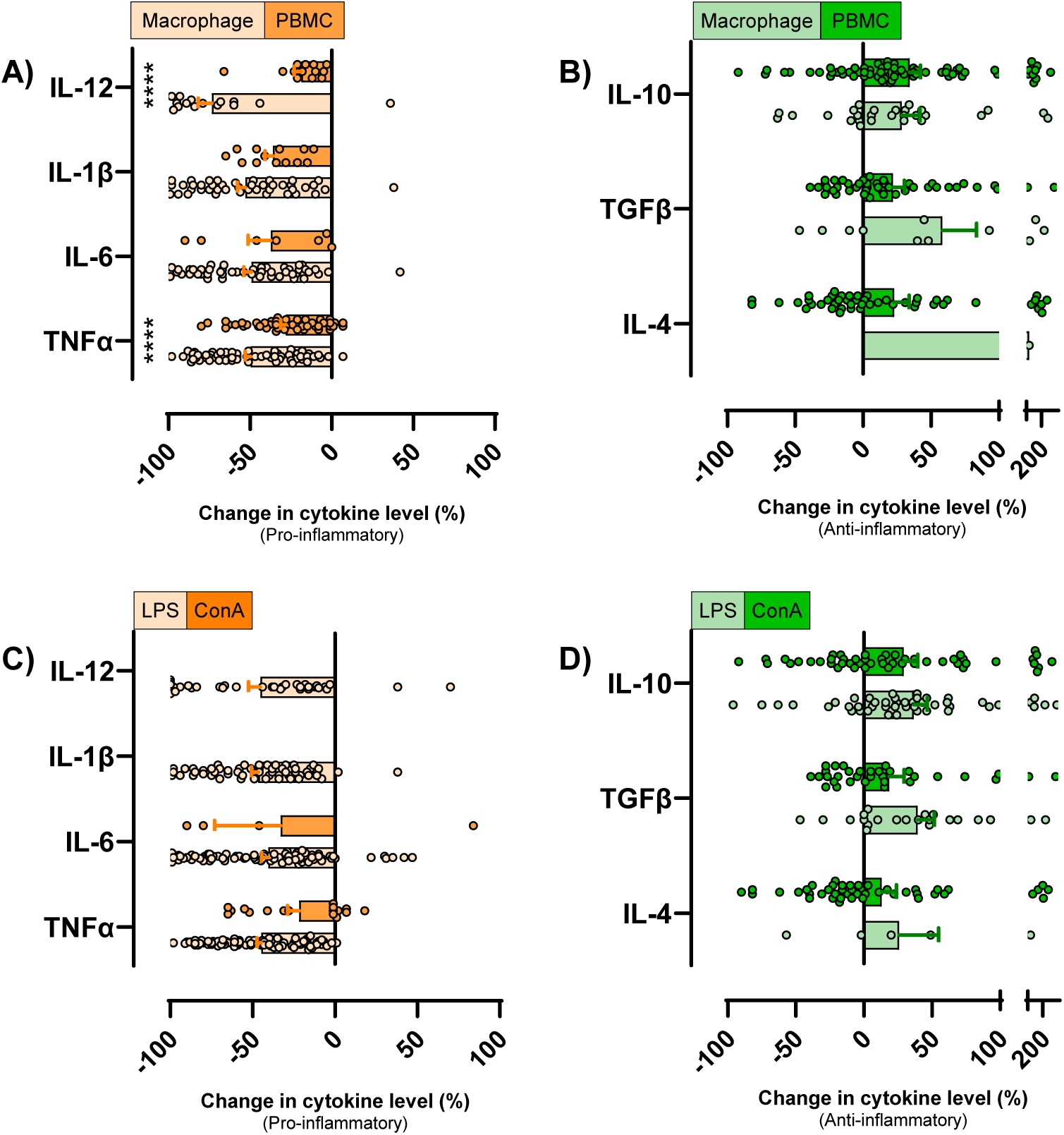
Effect of cell type and inflammagen on efficacy of HDPs. The effect of A & B) cell type and C & D) inflammagen on pro-inflammatory and anti-inflammatory cytokine levels. Each data point represents a specific record extracted from Table 1/Supplementary Excel file, and the mean±sem is also shown. Data were analyzed by two-way ANOVA with *post-hoc* Bonferroni. \*\*\*\**P*<0.0001 Macrophage vs. PBMC.

### Effects of HDPs from specific species on cytokines

The effects of the HDPs from the most widely used species were then assessed in terms of their effect on the most affected pro-inflammatory (IL-12, IL-1β, IL-6 and TNFα) and anti-inflammatory (IL-10, TGFβ and IL-4) cytokines (Fig. 5). This demonstrated that HDPs from all the widely used species significantly reduced pro-inflammatory cytokine levels (Fig. 5A; *Echinococcus granuloses: t*_(31)_=11.30, *P*<0.0001; *Schistosoma japonicum: t*_(16)_=9.18, *P*<0.0001; *Fasciola hepatica: t*_(47)_=9.03, *P*<0.0001; *Acanthocheilonema viteae: t*_(42)_=9.20, *P*<0.0001; *Schistosoma mansoni*: *t*_(20)_=7.41, *P*<0.001; *Haemonchus contortus: t*_(56)_=10.05, *P*<0.0001). There was less data available for the anti-inflammatory cytokines, but it was evident that HDPs from *Haemonchus contortus* were capable of both reducing pro-inflammatory cytokine levels, as well as significantly increasing anti-inflammatory cytokine levels (Fig. 5B; *Haemonchus contortus: t*_(139)_=5.61, *P*<0.0001).

**Fig. 5.**
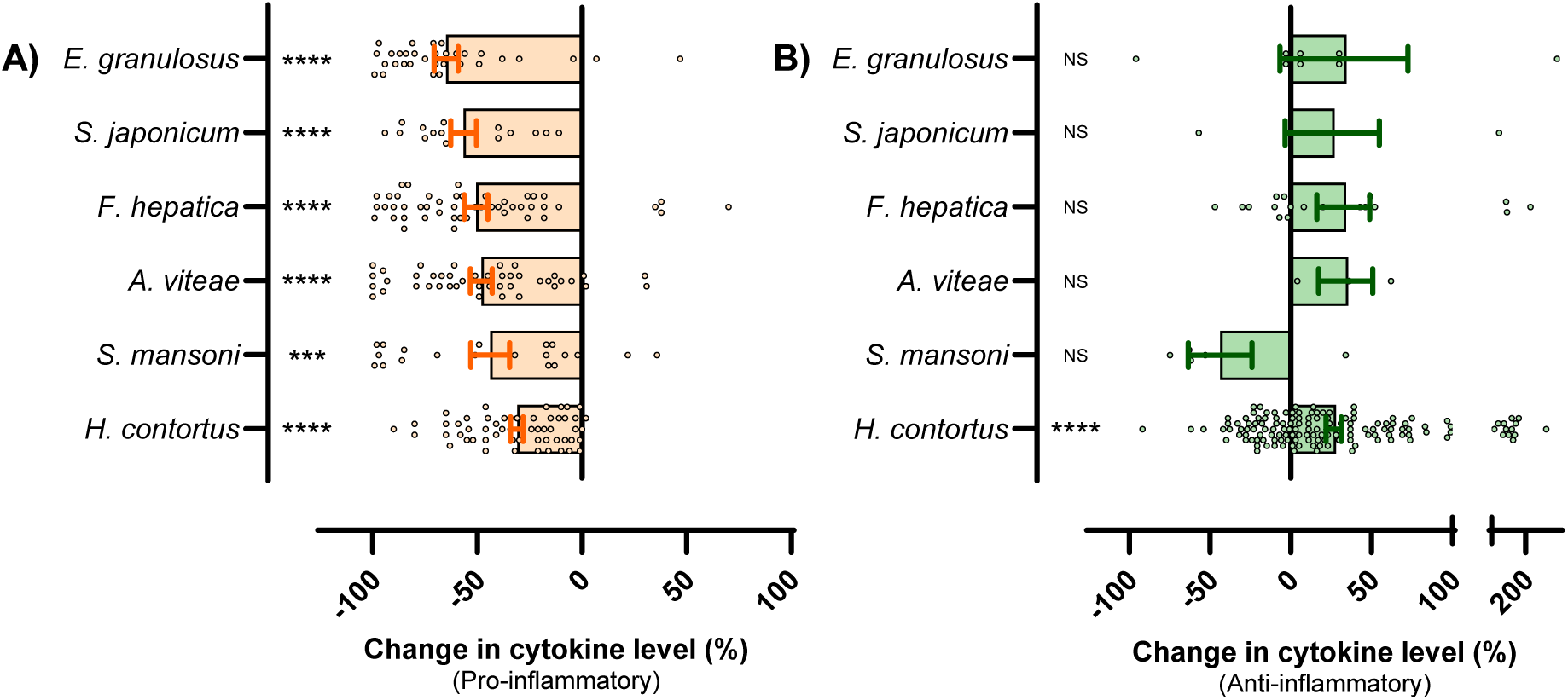
Effect of HDPs from specific species on cytokine levels. The effect of the HDPs from the most widely used species on A) pro-inflammatory (IL-12, IL-1β, IL-6 and TNFα) and b) anti-inflammatory (IL-10, TGFβ and IL-4) cytokine levels. Each data point represents a specific record extracted from Table 1/Supplementary Excel file, and the mean±sem is also shown. Data were analysed by one-sample test with the hypothesised population mean set to zero. \*\*\*\**P*<0.0001 and \*\*\**P*<0.001. NS: not-significantly different to zero.

### Effects of HDPs from specific species on specific cytokines

This data were then further subdivided to determine the effects of the HDPs from the most widely used species on the most affected pro-inflammatory cytokines individually (Fig. 6). This demonstrated that HDPs from *Echinococcus granuloses* significantly reduced levels of all of the most affected pro-inflammatory cytokines (Fig. 6: IL-12*: t*_(4)_=17.84, *P*<0.0001; IL-1β*: t*_(9)_=6.60, *P*<0.0001; IL-6*: t*_(9)_=5.14, *P*<0.001; TNFα*: t*_(6)_=5.21, *P*<0.01). HDPs from the other species also significantly reduced levels of most of the pro-inflammatory cytokines (statistical outcomes in Fig. 6).

**Fig. 6.**
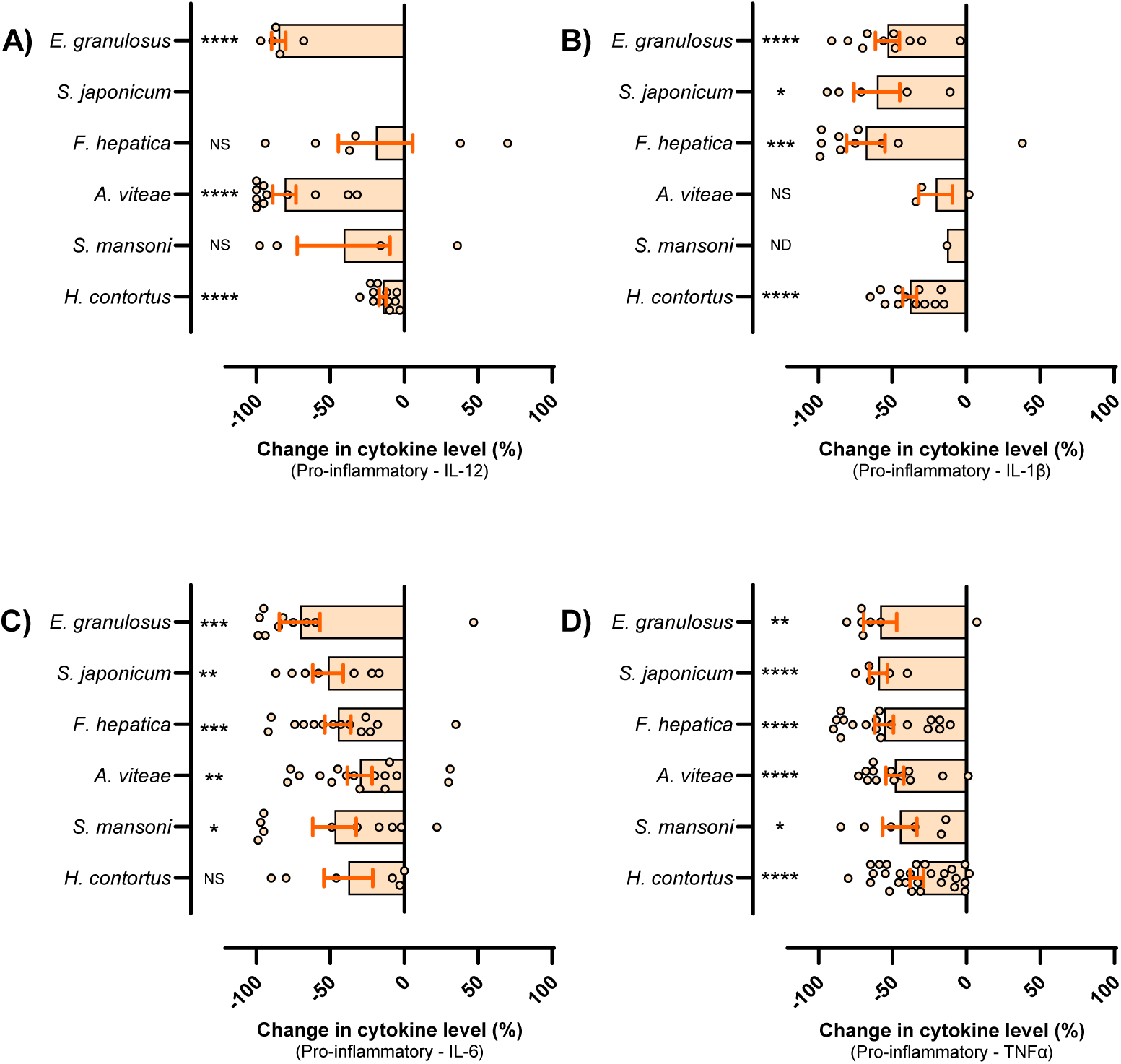
Effect of HDPs from specific species on pro-inflammatory cytokine levels. The effect of the HDPs from the most widely used species on A) IL-12, B) IL-1β, C) IL-6 and D) TNFα cytokine levels. Each data point represents a specific record extracted from Table 1/Supplementary Excel file, and the mean±sem is also shown. Data were analysed by one-sample test with the hypothesised population mean set to zero. \*\*\*\**P*<0.0001, \*\*\**P*<0.001, \*\**P*<0.01 and \**P*<0.05. NS: not-significantly different to zero. ND: not done as too few points.

Similarly, the data were also subdivided to determine the effects of the HDPs from the most widely used species on the most affected anti-inflammatory cytokines individually (Fig. 7). Although there was less data available for the anti-inflammatory cytokines, there were sufficient data points for *Haemonchus contortus* to note that HDPs from this species significantly increase IL-10 and (to a lesser extent) TGFβ levels, but do not significantly increase IL-4 levels (Fig. 7: IL-10*: t*_(57)_=4.91, *P*<0.0001; TGFβ*: t*_(40)_=2.58, P<0.05; IL-4*: t*_(40)_=1.94, P>0.05).

**Fig. 7.**
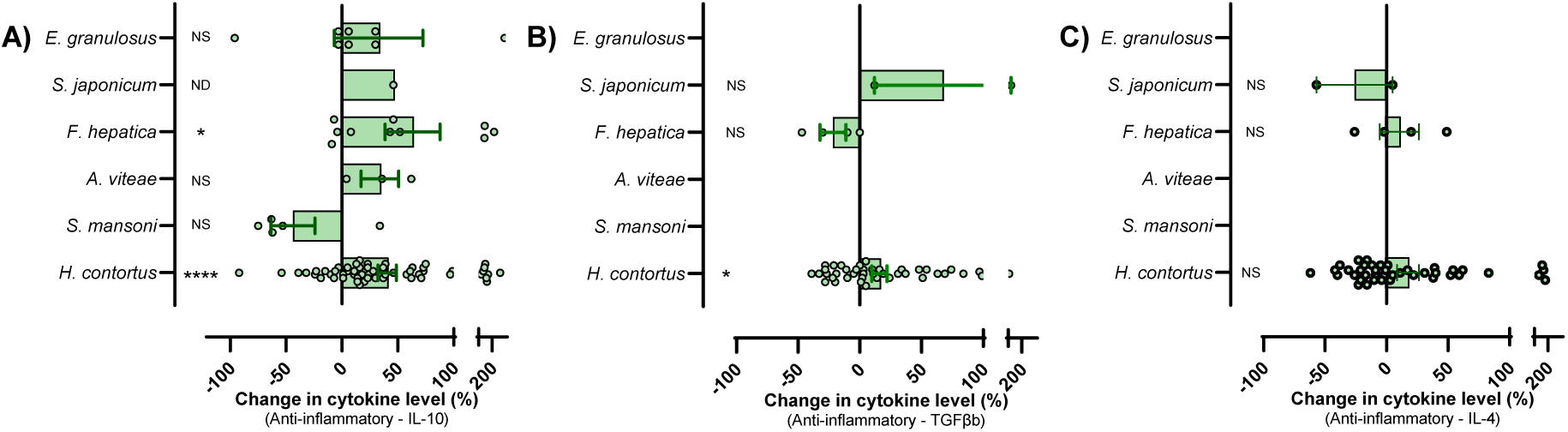
Effect of HDPs from specific species on anti-inflammatory cytokine levels. The effect of the HDPs from the most widely used species on A) IL-10, B) TGFβ and C) IL-4 cytokine levels. Each data point represents a specific record extracted from Table 1/Supplementary Excel file, and the mean±sem is also shown. Data were analysed by one-sample test with the hypothesised population mean set to zero. \*\*\*\**P*<0.0001 and \**P*<0.05. NS: not-significantly different to zero. ND: not done as too few points.

### Effects of mechanistic classes of HDPs on cytokines

The effect of HDPs was also assessed on the basis of their immunomodulatory mechanism of action for the main known mechanistic classes (Fig. 8). This demonstrated that HDPs that function as cysteine proteases were capable of both reducing pro-inflammatory (t_(99)_ = 13.58, P<0.0001) and increasing anti-inflammatory (t_(37)_ = 3.79, P<0.001) cytokine levels. In contrast, the other classes assessed (cathelicidin-like HDPs, ShK-related HDPs, fatty acid binding HDPs, glutathione transferase HDPs, galectin HPDs and thioredoxin peroxidase HDPs) were also capable of reducing pro-inflammatory cytokine levels but were less effective at increasing anti-inflammatory cytokine levels (or this was not assessed).

**Fig. 8.**
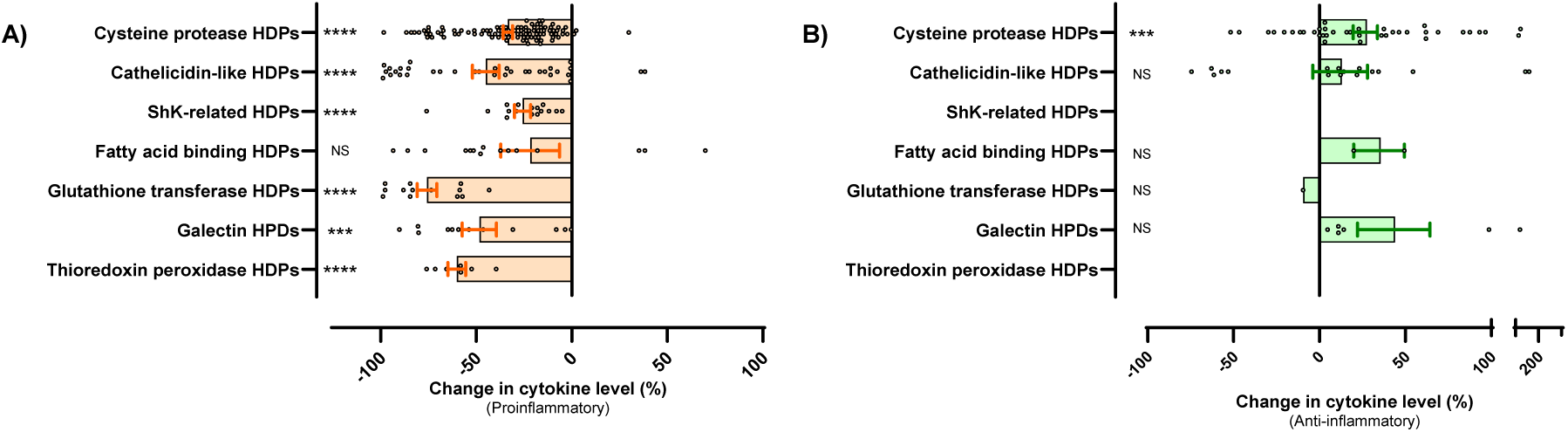
Effect of mechanistic classes of HDPs on cytokines. The effect of the HDPs from different mechanistic classes was assessed on A) pro-inflammatory (IL-12, IL-1β, IL-6 and TNFα) and B) anti-inflammatory (IL-10, TGFβ and IL-4) cytokine levels. Each data point represents a specific record extracted from Table 1/Supplementary Excel file, and the mean±sem is also shown. Data were analysed by one-sample test with the hypothesised population mean was set to zero. \*\*\*\**P*<0.0001 and \*\*\**P*<0.001.

### Effects of specific HDPs on cytokines

Finally, the data were plotted by individual HDP for the most widely assessed pro-inflammatory and anti-inflammatory cytokines (Fig. 9). The overwhelming trend for a reduction in proinflammatory cytokine levels (Fig. 9A) and an increase in anti-inflammatory cytokine levels (Fig. 9B) across HDPs is clear.

**Fig. 9.**
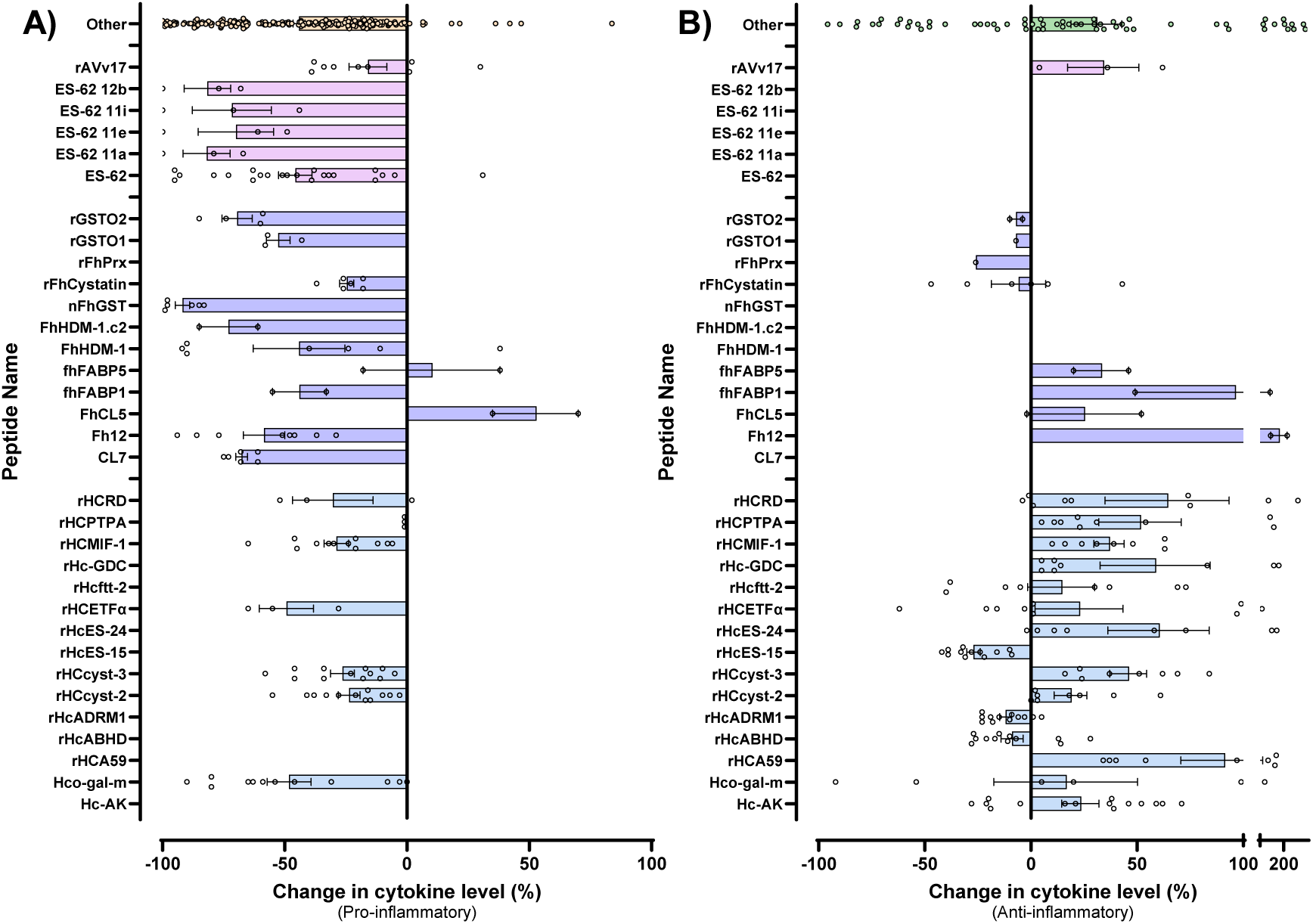
Effect of specific HDPs on cytokine levels. The effect of named HDPs from *H. contortus* (blue), *F. hepatica* (purple) and *A. viteae* (pink) on A) pro-inflammatory (IL-12, IL-1β, IL-6 and TNFα) and b) anti-inflammatory (IL-10, TGFβ and IL-4) cytokine levels. Each data point represents a specific record extracted from Table 1/Supplementary Excel file, and the mean±sem is also shown.

## Discussion

Secretory products of helminths have become a topic of interest in the last 20 years due to their immunomodulatory properties, leading researchers to investigate HPDs as therapeutic molecules for different immune and inflammatory conditions^1,4,10^. Preclinical studies in both cellular and animal models have assessed these immunomodulatory properties using various helminth species, HPDs, inflammagens, disease models, experimental parameters and measured outcomes in terms of functionality of immune cells and their secretome. In this review, we focussed on consolidating the studies in cellular models with a focus on the effect of HDPs on cytokine levels. Using a systematic approach, we identified 79 articles in which >60 HDPs from 20 helminth species were assessed largely in LPS or Concanavalin A stimulated macrophages, peripheral blood mononuclear cells or dendritic cells. Regardless of inflammagen, cell type or species, the overwhelming effect of the HDPs was a reduction in expression or production of pro-inflammatory cytokines such as IL-12, IL-1β, IL-6 and TNF, and an increase in anti-inflammatory cytokines such as IL-10, TGFβ and IL-4. Because of their profound effect on cytokine levels, this systematic review adds to the growing body of literature that supports the exploration of helminth secretory products as potential therapeutics in pathological conditions driven by overactive immune and inflammatory responses.

Although this review identified >60 HPDs from 20 helminth species, the most widely used species for derivation of HPDs were the nematodes, *Haemonchus contortus* and *Acanthocheilonema viteae,* the trematodes, *Fasciola hepatica*, *Schistosoma mansoni* and *Schistosoma japonicum*, and the cestode, *Echinococcus granuloses*. These have adapted to suppress their host’s immune system, allowing them to be tolerated by the host for significant periods of time. For example, the blood flukes, *Schistosoma mansoni* and *Schistosoma japonicum* have very long infection periods and can persist in humans for decades^91^, whereas, in contrast, infections by the liver fluke, *Fasciola hepatica*^92^, or the blood-sucking roundworm, *Haemonchus contortus*^93^, are typically shorter, lasting months or years. Regardless of the duration of the infective period, these endoparasites have evolved multiple mechanisms through which they can evade, modulate and suppress the immune system of the host species, including the secretion of immunomodulatory HDPs.

This study review revealed that >60 different HDPs have been assessed for their effect on cytokine responses in immune cells. The most common mechanistic class of the molecules assessed was cysteine protease activity which is associated with HDPs from many species^94–96^. Helminths, along with many other pathogens, have evolved to secrete cysteine proteases that facilitate invasion, infection and immune suppression of the host. These have various effects ranging from degradation (of extracellular matrix components, for example) of proteins to facilitate helminth invasion, to degradation of many proteins involved in the immune response. These cysteine protease HDPs can cleave the hinge region in IgG antibodies, cleave and inactivate pro-inflammatory cytokines, and degrade and inactivate pathogen-detecting Toll-like receptors among many other mechanisms^95^. This review demonstrated that HDPs with cysteine protease activity consistently and significantly reduced pro-inflammatory cytokine levels and increased anti-inflammatory cytokine levels, in line with the existing literature. Another mechanistic class that was widely assessed were the cathelicidin-like HPDs. These are the so-called helminth defense molecules (HDMs) secreted by trematodes such as *Fasciola hepatica*, *Schistosoma mansoni* and *Schistosoma japonicum*, that share that share structural and functional similarities with mammalian cathelicidin^97,98^. Like the cysteine proteases, these have multiple mechanisms through which they can enable trematode infection and immune suppression of the host including molecular mimicry, LPS-binding and sequestration, and inhibition of the NLRP3 inflammasome activation^99^. In the present review, the cathelicidin-like HDPs also profoundly reduced pro-inflammatory cytokine levels but effects on anti-inflammatory cytokines were not significant overall. Similarly, the other mechanistic classes assessed included ShK-related HDPs, fatty acid binding HDPs, glutathione transferase HDPs, galectin HPDs, and thioredoxin peroxidase HDPs all of which, like the cathelicidin-like HDPs, significantly reduced proinflammatory cytokine levels.

One unexpected aspect of the data consolidation in this review was the finding of a greater efficacy of HDPs (greater suppression of pro-inflammatory cytokines) in macrophages compared with PBMCs. However, this is likely simply due to the heterogeneity of the PBMC population which includes many populations of lymphocytes (including T-cells, B-cells and natural killer cells) as well as monocytes which can be considered immature macrophages. Thus, relative to more uniform macrophage cultures, PBMC cultures are likely to have variable responsivity to the main inflammagens used and consequently, variability responsivity to the HDPs. Another interesting aspect of the current systematic review is that it revealed the paucity of studies using other, critically important, immune cells. For example, there was only a single study each in which the immunomodulatory effect of HDPs in mast cells^49^ or microglia^32^ was assessed. This highlights an important gap in literature as there is potential for the HPDs to be effective immunomodulators in allergic and/or neuroinflammatory conditions. Another gap in the literature that was identified in this systematic review was the lack of variability in the inflammagen used with most studies using LPS. LPS is a gram-negative bacterial endotoxin that is recognized by Toll-like receptor 4 (TLR4) on multiple cell types, causing a signaling cascade which leads to the activation of NF-kB, triggering the release of proinflammatory molecules like TNFα, IL-6, and nitric oxide^100^. Although LPS-induced inflammation is a widely accepted and common model for many inflammatory conditions, there is a need to test whether HDPs can mitigate the effects of other clinically relevant inflammatory triggers, not only other pathogen-associated molecular patterns, but also damage-associated molecular patterns.

While this review focused on assessing the efficacy of HDPs (in modulating cytokine levels) in cellular models, several studies have also assessed their efficacy in animal models on inflammatory disease such as inflammatory bowel disease^4^, multiple sclerosis^90^, rheumatoid arthritis^101^ and asthma^102^. The success of these, and other studies, studies led to a series of clinical trials of helminth therapy in which patients are infected with specific helminths to modulate the immune system and treat their inflammatory and autoimmune diseases. Infection with the hookworm, *Necator americanus*, for example, has been assessed in clinical trials of ulcerative colitis^103^, multiple sclerosis^104^ and celiac disease^105^ with mixed results in terms of both efficacy and safety. This highlights the rationale for continued research and development of HDPs as therapeutic drugs as these would reduce the risks associated with live parasitic infection. To date, only one clinical trial of a HDP has been completed (NCT02281916) in which P28GST, a *Schistosoma haematobium*-derived glutathione S-transferase, was tested in patients with Crohn’s disease with some evidence of efficacy^106^.

Overall, this review systematically consolidates and summarises the effect of immunomodulatory HDPs in cellular model of inflammation. It clearly demonstrates that suppression of pro-inflammatory, and enhancement of anti-inflammatory, cytokine expression and production in response to inflammagen stimulation in immune cells is a property of many HDPs from many species. The review also revealed important gaps in this literature highlighting the opportunity to explore the immunomodulatory efficacy of HDPs in other cell types using other inflammatory stimuli to determine their potential for other immune system disorders heretofore relatively unexplored. Consolidating this literature may direct future research into the use of HDPs in a wider variety of preclinical models in order to determine their broader therapeutic potential.

## Acknowledgements

SS is funded by the Parkinson’s Disease Research Award from Tony & Peigí O’Donoghue through the Galway University Foundation. ED would also like to acknowledge grants from the Michael J Fox Foundation for Parkinson’s Research (Grant Numbers: 17244 and 023410).

## Data availability

All data is available in the Supplementary Excel file.

## Author Contributions

SS, AF and ED led the design and implementation of the systematic review, and wrote the first draft of the manuscript; RL, SD, JPD and DMK provided expert input into the drafts of the manuscript.

## REFERENCES

1 Yeshi, K., Ruscher, R., Loukas, A. & Wangchuk, P. Immunomodulatory and biological properties of helminth-derived small molecules: Potential applications in diagnostics and therapeutics. Front Parasitol 1, 984152, doi:10.3389/fpara.2022.984152 (2022).

2 Chen, J., Gong, Y., Chen, Q., Li, S. & Zhou, Y. Global burden of soil-transmitted helminth infections, 1990-2021. Infect Dis Poverty 13, 77, doi:10.1186/s40249-024-01238-9 (2024).

3 Caldrer, S., Ursini, T., Santucci, B., Motta, L. & Angheben, A. Soil-Transmitted Helminths and Anaemia: A Neglected Association Outside the Tropics. Microorganisms 10, doi:10.3390/microorganisms10051027 (2022).

4 Arai, T. & Lopes, F. Potential of human helminth therapy for resolution of inflammatory bowel disease: The future ahead. Exp Parasitol 232, 108189, doi:10.1016/j.exppara.2021.108189 (2022).

5 Szuba, M. et al. Geohelminths: Use in the Treatment of Selected Human Diseases. Pathogens 13, doi:10.3390/pathogens13080703 (2024).

6 Versini, M. et al. Unraveling the Hygiene Hypothesis of helminthes and autoimmunity: origins, pathophysiology, and clinical applications. BMC Med 13, 81, doi:10.1186/s12916-015-0306-7 (2015).

7 Mu, Y., McManus, D. P., Hou, N. & Cai, P. Schistosome Infection and Schistosome-Derived Products as Modulators for the Prevention and Alleviation of Immunological Disorders. Front Immunol 12, 619776, doi:10.3389/fimmu.2021.619776 (2021).

8 Fumagalli, M. et al. The landscape of human genes involved in the immune response to parasitic worms. BMC Evol Biol 10, 264, doi:10.1186/1471-2148-10-264 (2010).

9 Kasal, D. N., Warner, L. M., Bryant, A. S., Tait Wojno, E. & von Moltke, J. Systemic Immune Modulation by Gastrointestinal Nematodes. Annu Rev Immunol 42, 259–288, doi:10.1146/annurev-immunol-090222-101331 (2024).

10 Alghanmi, M. et al. Helminth-derived proteins as immune system regulators: a systematic review of their promise in alleviating colitis. BMC Immunol 25, 21, doi:10.1186/s12865-024-00614-2 (2024).

11 Page, M. J. et al. The PRISMA 2020 statement: an updated guideline for reporting systematic reviews. BMJ 372, n71, doi:10.1136/bmj.n71 (2021).

12 Schonemeyer, A. et al. Modulation of human T cell responses and macrophage functions by onchocystatin, a secreted protein of the filarial nematode Onchocerca volvulus. J Immunol 167, 3207–3215, doi:10.4049/jimmunol.167.6.3207 (2001).

13 Goodridge, H. S. et al. Modulation of macrophage cytokine production by ES-62, a secreted product of the filarial nematode Acanthocheilonema viteae. J Immunol 167, 940–945, doi:10.4049/jimmunol.167.2.940 (2001).

14 Spolski, R. J., Thomas, P. G., See, E. J., Mooney, K. A. & Kuhn, R. E. Larval Taenia crassiceps secretes a protein with characteristics of murine interferon-gamma. Parasitol Res 88, 431–438, doi:10.1007/s00436-002-0590-y (2002).

15 McInnes, I. B. et al. A novel therapeutic approach targeting articular inflammation using the filarial nematode-derived phosphorylcholine-containing glycoprotein ES-62. J Immunol 171, 2127–2133, doi:10.4049/jimmunol.171.4.2127 (2003).

16 Goodridge, H. S. et al. Immunomodulation via novel use of TLR4 by the filarial nematode phosphorylcholine-containing secreted product, ES-62. J Immunol 174, 284–293, doi:10.4049/jimmunol.174.1.284 (2005).

17 Rigano, R. et al. Echinococcus granulosus antigen B impairs human dendritic cell differentiation and polarizes immature dendritic cell maturation towards a Th2 cell response. Infect Immun 75, 1667–1678, doi:10.1128/IAI.01156-06 (2007).

18 Donnelly, S. et al. Helminth 2-Cys peroxiredoxin drives Th2 responses through a mechanism involving alternatively activated macrophages. FASEB J 22, 4022–4032, doi:10.1096/fj.08-106278 (2008).

19 Brannstrom, K., Sellin, M. E., Holmfeldt, P., Brattsand, M. & Gullberg, M. The Schistosoma mansoni protein Sm16/SmSLP/SmSPO-1 assembles into a nine-subunit oligomer with potential To inhibit Toll-like receptor signaling. Infect Immun 77, 1144–1154, doi:10.1128/IAI.01126-08 (2009).

20 Sun, X. J. et al. Unique roles of Schistosoma japonicum protein Sj16 to induce IFN-gamma and IL-10 producing CD4(+)CD25(+) regulatory T cells in vitro and in vivo. Parasite Immunol 34, 430–439, doi:10.1111/j.1365-3024.2012.01377.x (2012).

21 Chhabra, S. et al. Kv1.3 channel-blocking immunomodulatory peptides from parasitic worms: implications for autoimmune diseases. FASEB J 28, 3952–3964, doi:10.1096/fj.14-251967 (2014).

22 Wang, W. et al. Galectin Hco-gal-m from Haemonchus contortus modulates goat monocytes and T cell function in different patterns. Parasit Vectors 7, 342, doi:10.1186/1756-3305-7-342 (2014).

23 Du, L. et al. Regulation of recombinant Trichinella spiralis 53-kDa protein (rTsP53) on alternatively activated macrophages via STAT6 but not IL-4Ralpha in vitro. Cell Immunol 288, 1–7, doi:10.1016/j.cellimm.2014.01.010 (2014).

24 Shen, J. et al. Gene expression profile of LPS-stimulated dendritic cells induced by a recombinant Sj16 (rSj16) derived from Schistosoma japonicum. Parasitol Res 113, 3073–3083, doi:10.1007/s00436-014-3973-y (2014).

25 Figueroa-Santiago, O. & Espino, A. M. Fasciola hepatica fatty acid binding protein induces the alternative activation of human macrophages. Infect Immun 82, 5005–5012, doi:10.1128/IAI.02541-14 (2014).

26 Dlugosz, E., Wasyl, K., Klockiewicz, M. & Wisniewski, M. Toxocara canis mucins among other excretory-secretory antigens induce in vitro secretion of cytokines by mouse splenocytes. Parasitol Res 114, 3365–3371, doi:10.1007/s00436-015-4561-5 (2015).

27 Ferguson, B. J. et al. The Schistosoma mansoni T2 ribonuclease omega-1 modulates inflammasome-dependent IL-1beta secretion in macrophages. Int J Parasitol 45, 809–813, doi:10.1016/j.ijpara.2015.08.005 (2015).

28 Wang, Y., Wang, Q., Lv, S. & Zhang, S. Different protein of Echinococcus granulosus stimulates dendritic induced immune response. Parasitology 142, 879–889, doi:10.1017/S0031182014002005 (2015).

29 Sanin, D. E. & Mountford, A. P. Sm16, a major component of Schistosoma mansoni cercarial excretory/secretory products, prevents macrophage classical activation and delays antigen processing. Parasit Vectors 8, 1, doi:10.1186/s13071-014-0608-1 (2015).

30 Martin, I., Caban-Hernandez, K., Figueroa-Santiago, O. & Espino, A. M. Fasciola hepatica fatty acid binding protein inhibits TLR4 activation and suppresses the inflammatory cytokines induced by lipopolysaccharide in vitro and in vivo. J Immunol 194, 3924–3936, doi:10.4049/jimmunol.1401182 (2015).

31 Khatri, V., Amdare, N., Tarnekar, A., Goswami, K. & Reddy, M. V. Brugia malayi cystatin therapeutically ameliorates dextran sulfate sodium-induced colitis in mice. J Dig Dis 16, 585–594, doi:10.1111/1751-2980.12290 (2015).

32 Behrendt, P. et al. A Helminth Protease Inhibitor Modulates the Lipopolysaccharide-Induced Proinflammatory Phenotype of Microglia in vitro. Neuroimmunomodulation 23, 109–121, doi:10.1159/000444756 (2016).

33 Gadahi, J. A. et al. Recombinant Haemonchus contortus 24 kDa excretory/secretory protein (rHcES-24) modulate the immune functions of goat PBMCs in vitro. Oncotarget 7, 83926–83937, doi:10.18632/oncotarget.13487 (2016).

34 Cao, X. et al. iTRAQ-based comparative proteomic analysis of excretory-secretory proteins of schistosomula and adult worms of Schistosoma japonicum. J Proteomics 138, 30–39, doi:10.1016/j.jprot.2016.02.015 (2016).

35 Gadahi, J. A. et al. Recombinant protein of Haemonchus contortus 14-3-3 isoform 2 (rHcftt-2) decreased the production of IL-4 and suppressed the proliferation of goat PBMCs in vitro. Exp Parasitol 171, 57–66, doi:10.1016/j.exppara.2016.10.014 (2016).

36 Eason, R. J. et al. The helminth product, ES-62 modulates dendritic cell responses by inducing the selective autophagolysosomal degradation of TLR-transducers, as exemplified by PKCdelta. Sci Rep 6, 37276, doi:10.1038/srep37276 (2016).

37 Silva-Alvarez, V. et al. Echinococcus granulosus Antigen B binds to monocytes and macrophages modulating cell response to inflammation. Parasit Vectors 9, 69, doi:10.1186/s13071-016-1350-7 (2016).

38 Lund, M. E. et al. A parasite-derived 68-mer peptide ameliorates autoimmune disease in murine models of Type 1 diabetes and multiple sclerosis. Sci Rep 6, 37789, doi:10.1038/srep37789 (2016).

39 Alvarado, R. et al. The immune modulatory peptide FhHDM-1 secreted by the helminth Fasciola hepatica prevents NLRP3 inflammasome activation by inhibiting endolysosomal acidification in macrophages. FASEB J 31, 85–95, doi:10.1096/fj.201500093R (2017).

40 Ehsan, M. et al. Arginine kinase from Haemonchus contortus decreased the proliferation and increased the apoptosis of goat PBMCs in vitro. Parasit Vectors 10, 311, doi:10.1186/s13071-017-2244-z (2017).

41 Floudas, A. et al. Composition of the Schistosoma mansoni worm secretome: Identification of immune modulatory Cyclophilin A. PLoS Negl Trop Dis 11, e0006012, doi:10.1371/journal.pntd.0006012 (2017).

42 Wang, Y. et al. Characterization of a secreted macrophage migration inhibitory factor homologue of the parasitic nematode Haemonchus Contortus acting at the parasite-host cell interface. Oncotarget 8, 40052–40064, doi:10.18632/oncotarget.16675 (2017).

43 Wang, Y. et al. Characterization of a secreted cystatin of the parasitic nematode Haemonchus contortus and its immune-modulatory effect on goat monocytes. Parasit Vectors 10, 425, doi:10.1186/s13071-017-2368-1 (2017).

44 Wang, Y. et al. Modulation of goat monocyte function by HCcyst-2, a secreted cystatin from Haemonchus contortus. Oncotarget 8, 44108–44120, doi:10.18632/oncotarget.17308 (2017).

45 Lumb, F. E. et al. Dendritic cells provide a therapeutic target for synthetic small molecule analogues of the parasitic worm product, ES-62. Sci Rep 7, 1704, doi:10.1038/s41598-017-01651-1 (2017).

46 Amdare, N. P. et al. Therapeutic potential of the immunomodulatory proteins Wuchereria bancrofti L2 and Brugia malayi abundant larval transcript 2 against streptozotocin-induced type 1 diabetes in mice. J Helminthol 91, 539–548, doi:10.1017/S0022149X1600064X (2017).

47 Wang, X. et al. Inhibition of cytokine response to TLR stimulation and alleviation of collagen-induced arthritis in mice by Schistosoma japonicum peptide SJMHE1. J Cell Mol Med 21, 475–486, doi:10.1111/jcmm.12991 (2017).

48 Knuhr, K. et al. Schistosoma mansoni Egg-Released IPSE/alpha-1 Dampens Inflammatory Cytokine Responses via Basophil Interleukin (IL)-4 and IL-13. Front Immunol 9, 2293, doi:10.3389/fimmu.2018.02293 (2018).

49 Ball, D. H., Al-Riyami, L., Harnett, W. & Harnett, M. M. IL-33/ST2 signalling and crosstalk with FcepsilonRI and TLR4 is targeted by the parasitic worm product, ES-62. Sci Rep 8, 4497, doi:10.1038/s41598-018-22716-9 (2018).

50 Zheng, Y. et al. Identification of emu-TegP11, an EF-hand domain-containing tegumental protein of Echinococcus multilocularis. Vet Parasitol 255, 107–113, doi:10.1016/j.vetpar.2018.04.006 (2018).

51 Togre, N. et al. Immunomodulatory potential of recombinant filarial protein, rWbL2, and its therapeutic implication in experimental ulcerative colitis in mouse. Immunopharmacol Immunotoxicol 40, 483–490, doi:10.1080/08923973.2018.1431925 (2018).

52 Wang, H. et al. Thioredoxin peroxidase secreted by Echinococcus granulosus (sensu stricto) promotes the alternative activation of macrophages via PI3K/AKT/mTOR pathway. Parasit Vectors 12, 542, doi:10.1186/s13071-019-3786-z (2019).

53 Aguayo, V. et al. Fasciola hepatica GST downregulates NF-kappaB pathway effectors and inflammatory cytokines while promoting survival in a mouse septic shock model. Sci Rep 9, 2275, doi:10.1038/s41598-018-37652-x (2019).

54 Wang, Q. et al. Hepatocellular carcinoma-associated antigen 59 of Haemonchus contortus modulates the functions of PBMCs and the differentiation and maturation of monocyte-derived dendritic cells of goats in vitro. Parasit Vectors 12, 105, doi:10.1186/s13071-019-3375-1 (2019).

55 Vacca, F. et al. A helminth-derived suppressor of ST2 blocks allergic responses. Elife 9, doi:10.7554/eLife.54017 (2020).

56 Ehsan, M. et al. Characterization of Haemonchus contortus Excretory/Secretory Antigen (ES-15) and Its Modulatory Functions on Goat Immune Cells In Vitro. Pathogens 9, doi:10.3390/pathogens9030162 (2020).

57 Ilgova, J., Kavanova, L., Matiaskova, K., Salat, J. & Kasny, M. Effect of cysteine peptidase inhibitor of Eudiplozoon nipponicum (Monogenea) on cytokine expression of macrophages in vitro. Mol Biochem Parasitol 235, 111248, doi:10.1016/j.molbiopara.2019.111248 (2020).

58 Kobpornchai, P. et al. A novel cystatin derived from Trichinella spiralis suppresses macrophage-mediated inflammatory responses. PLoS Negl Trop Dis 14, e0008192, doi:10.1371/journal.pntd.0008192 (2020).

59 Shiels, J. et al. Schistosoma mansoni immunomodulatory molecule Sm16/SPO-1/SmSLP is a member of the trematode-specific helminth defence molecules (HDMs). PLoS Negl Trop Dis 14, e0008470, doi:10.1371/journal.pntd.0008470 (2020).

60 Wang, Y. et al. Characterization of a rhodanese homologue from Haemonchus contortus and its immune-modulatory effects on goat immune cells in vitro. Parasit Vectors 13, 454, doi:10.1186/s13071-020-04333-6 (2020).

61 Wang, Y. et al. Characterization of a phosphotyrosyl phosphatase activator homologue of the parasitic nematode Haemonchus contortus and its immunomodulatory effect on goat peripheral blood mononuclear cells in vitro. Int J Parasitol 50, 1157–1166, doi:10.1016/j.ijpara.2020.07.004 (2020).

62 Wang, Y. et al. Modulatory functions of recombinant electron transfer flavoprotein alpha subunit protein from Haemonchus contortus on goat immune cells in vitro. Vet Parasitol 288, 109300, doi:10.1016/j.vetpar.2020.109300 (2020).

63 Naqvi, M. A. et al. Galectin Domain Containing Protein from Haemonchus contortus Modulates the Immune Functions of Goat PBMCs and Regulates CD4+ T-Helper Cells In Vitro. Biomolecules 10, doi:10.3390/biom10010116 (2020).

64 Lu, M. et al. Unveiling the immunomodulatory properties of Haemonchus contortus adhesion regulating molecule 1 interacting with goat T cells. Parasit Vectors 13, 424, doi:10.1186/s13071-020-04297-7 (2020).

65 Lu, M. et al. A Novel alpha/beta Hydrolase Domain Protein Derived From Haemonchus contortus Acts at the Parasite-Host Interface. Front Immunol 11, 1388, doi:10.3389/fimmu.2020.01388 (2020).

66 Montero, L., Cervantes-Torres, J., Sciutto, E. & Fragoso, G. Helminth-derived peptide GK-1 induces Myd88-dependent pro-inflammatory signaling events in bone marrow-derived antigen-presenting cells. Mol Immunol 128, 22–32, doi:10.1016/j.molimm.2020.09.015 (2020).

67 Buitrago, G. et al. A netrin domain-containing protein secreted by the human hookworm Necator americanus protects against CD4 T cell transfer colitis. Transl Res 232, 88–102, doi:10.1016/j.trsl.2021.02.012 (2021).

68 Ehsan, M. et al. Fasciola gigantica tegumental calcium-binding EF-hand protein 4 exerts immunomodulatory effects on goat monocytes. Parasit Vectors 14, 276, doi:10.1186/s13071-021-04784-5 (2021).

69 Xie, H. et al. Schistosoma japonicum Cystatin Alleviates Sepsis Through Activating Regulatory Macrophages. Front Cell Infect Microbiol 11, 617461, doi:10.3389/fcimb.2021.617461 (2021).

70. Corbet, M. et al. Suppression of inflammatory arthritis by the parasitic worm product ES-62 is associated with epigenetic changes in synovial fibroblasts. PLoS Pathog 17, e1010069, doi:10.1371/journal.ppat.1010069 (2021).

71 Smallwood, T. B. et al. Synthetic hookworm-derived peptides are potent modulators of primary human immune cell function that protect against experimental colitis in vivo. J Biol Chem 297, 100834, doi:10.1016/j.jbc.2021.100834 (2021).

72 Ryan, S. M. et al. Novel antiinflammatory biologics shaped by parasite-host coevolution. Proc Natl Acad Sci U S A 119, e2202795119, doi:10.1073/pnas.2202795119 (2022).

73 Kobpornchai, P., Reamtong, O., Phuphisut, O., Malaitong, P. & Adisakwattana, P. Serine protease inhibitor derived from Trichinella spiralis (TsSERP) inhibits neutrophil elastase and impairs human neutrophil functions. Front Cell Infect Microbiol 12, 919835, doi:10.3389/fcimb.2022.919835 (2022).

74 Wang, X. et al. Molecular characterization of a novel GSTO2 of Fasciola hepatica and its roles in modulating murine macrophages. Parasite 29, 16, doi:10.1051/parasite/2022016 (2022).

75 Loghry, H. J., Sondjaja, N. A., Minkler, S. J. & Kimber, M. J. Secreted filarial nematode galectins modulate host immune cells. Front Immunol 13, 952104, doi:10.3389/fimmu.2022.952104 (2022).

76 Zhang, K. et al. Molecular Characteristics and Potent Immunomodulatory Activity of Fasciola hepatica Cystatin. Korean J Parasitol 60, 117–126, doi:10.3347/kjp.2022.60.2.117 (2022).

77 Li, H. et al. Trichinella spiralis cystatin alleviates polymicrobial sepsis through activating regulatory macrophages. Int Immunopharmacol 109, 108907, doi:10.1016/j.intimp.2022.108907 (2022).

78 Zawistowska-Deniziak, A. et al. Fasciola hepatica Fatty Acid Binding Protein 1 Modulates T cell Polarization by Promoting Dendritic Cell Thrombospondin-1 Secretion Without Affecting Metabolic Homeostasis in Obese Mice. Front Immunol 13, 884663, doi:10.3389/fimmu.2022.884663 (2022).

79 Ruiz-Jimenez, C. et al. Fasciola hepatica fatty acid binding protein (Fh12) induces apoptosis and tolerogenic properties in murine bone marrow derived dendritic cells. Exp Parasitol 231, 108174, doi:10.1016/j.exppara.2021.108174 (2021).

80 Stachyra, A. & Wesolowska, A. Immunomodulatory in vitro effects of Trichinella cystatin-like protein on mouse splenocytes. Exp Parasitol 252, 108585, doi:10.1016/j.exppara.2023.108585 (2023).

81 Xifeng, W. et al. The regulatory roles of Fasciola hepatica GSTO1 protein in inflammatory cytokine expression and apoptosis in murine macrophages. Acta Trop 245, 106977, doi:10.1016/j.actatropica.2023.106977 (2023).

82 Chantree, P. et al. Type I Cystatin Derived from Fasciola gigantica Suppresses Macrophage-Mediated Inflammatory Responses. Pathogens 12, doi:10.3390/pathogens12030395 (2023).

83 Quinteros, S. L. et al. The helminth derived peptide FhHDM-1 redirects macrophage metabolism towards glutaminolysis to regulate the pro-inflammatory response. Front Immunol 14, 1018076, doi:10.3389/fimmu.2023.1018076 (2023).

84 Colomb, F. et al. IL-33-binding HpARI family homologues with divergent effects in suppressing or enhancing type 2 immune responses. Infect Immun 92, e0039523, doi:10.1128/iai.00395-23 (2024).

85 Yang, Q. et al. Type I Cystatin Derived from Cysticercus pisiformis-Stefins, Suppresses LPS-Mediated Inflammatory Response in RAW264.7 Cells. Microorganisms 12, doi:10.3390/microorganisms12050850 (2024).

86 Ehsan, M. et al. Immune modulation of goat monocytes by Fasciola gigantica Legumain-1 protein (Fg-LGMN-1). Exp Parasitol 256, 108671, doi:10.1016/j.exppara.2023.108671 (2024).

87 Folle, A. M. et al. Modulatory actions of Echinococcus granulosus antigen B on macrophage inflammatory activation. Front Cell Infect Microbiol 14, 1362765, doi:10.3389/fcimb.2024.1362765 (2024).

88 Gong, J. Z. et al. Cathepsin L of Fasciola hepatica meliorates colitis by altering the gut microbiome and inflammatory macrophages. Int J Biol Macromol 286, 138270, doi:10.1016/j.ijbiomac.2024.138270 (2025).

89 Li, X. et al. Echinococcus multilocularis serpin regulates macrophage polarization and reduces gut dysbiosis in colitis. Infect Immun 92, e0023224, doi:10.1128/iai.00232-24 (2024).

90 Lalor, R. et al. An immunoregulatory amphipathic peptide derived from Fasciola hepatica helminth defense molecule (FhHDM-1.C2) exhibits potent biotherapeutic activity in a murine model of multiple sclerosis. FASEB J 39, e70380, doi:10.1096/fj.202400793RR (2025).

91 Lo, N. C. et al. Review of 2022 WHO guidelines on the control and elimination of schistosomiasis. Lancet Infect Dis 22, e327–e335, doi:10.1016/S1473-3099(22)00221-3 (2022).

92 Moazeni, M. & Ahmadi, A. Controversial aspects of the life cycle of Fasciola hepatica. Exp Parasitol 169, 81–89, doi:10.1016/j.exppara.2016.07.010 (2016).

93 Gasser, R. B., Schwarz, E. M., Korhonen, P. K. & Young, N. D. Understanding Haemonchus contortus Better Through Genomics and Transcriptomics. Adv Parasitol 93, 519–567, doi:10.1016/bs.apar.2016.02.015 (2016).

94 Robinson, M. W., Dalton, J. P. & Donnelly, S. Helminth pathogen cathepsin proteases: it’s a family affair. Trends Biochem Sci 33, 601–608, doi:10.1016/j.tibs.2008.09.001 (2008).

95 Donnelly, S., Dalton, J. P. & Robinson, M. W. How pathogen-derived cysteine proteases modulate host immune responses. Adv Exp Med Biol 712, 192–207, doi:10.1007/978-1-4419-8414-2_12 (2011).

96 Khatri, V., Chauhan, N. & Kalyanasundaram, R. Parasite Cystatin: Immunomodulatory Molecule with Therapeutic Activity against Immune Mediated Disorders. Pathogens 9, doi:10.3390/pathogens9060431 (2020).

97 Alvarado, R., O’Brien, B., Tanaka, A., Dalton, J. P. & Donnelly, S. A parasitic helminth-derived peptide that targets the macrophage lysosome is a novel therapeutic option for autoimmune disease. Immunobiology 220, 262–269, doi:10.1016/j.imbio.2014.11.008 (2015).

98 Ryan, S., Shiels, J., Taggart, C. C., Dalton, J. P. & Weldon, S. Fasciola hepatica-Derived Molecules as Regulators of the Host Immune Response. Front Immunol 11, 2182, doi:10.3389/fimmu.2020.02182 (2020).

99 Maizels, R. M., Smits, H. H. & McSorley, H. J. Modulation of Host Immunity by Helminths: The Expanding Repertoire of Parasite Effector Molecules. Immunity 49, 801–818, doi:10.1016/j.immuni.2018.10.016 (2018).

100 Sasai, M. & Yamamoto, M. Pathogen recognition receptors: ligands and signaling pathways by Toll-like receptors. Int Rev Immunol 32, 116–133, doi:10.3109/08830185.2013.774391 (2013).

101 Langdon, K. et al. Helminth-based therapies for rheumatoid arthritis: A systematic review and meta-analysis. Int Immunopharmacol 66, 366–372, doi:10.1016/j.intimp.2018.11.034 (2019).

102 Fernandes, J. S., Cardoso, L. S., Pitrez, P. M. & Cruz, A. A. Helminths and Asthma: Risk and Protection. Immunol Allergy Clin North Am 39, 417–427, doi:10.1016/j.iac.2019.03.009 (2019).

103 Mules, T. C. et al. Controlled Hookworm Infection for Medication-free Maintenance in Patients with Ulcerative Colitis: A Pilot, Double-blind, Randomized Control Trial. Inflamm Bowel Dis 30, 735–745, doi:10.1093/ibd/izad110 (2024).

104 Tanasescu, R. et al. Hookworm Treatment for Relapsing Multiple Sclerosis: A Randomized Double-Blinded Placebo-Controlled Trial. JAMA Neurol 77, 1089–1098, doi:10.1001/jamaneurol.2020.1118 (2020).

105 Croese, J. et al. Randomized, Placebo Controlled Trial of Experimental Hookworm Infection for Improving Gluten Tolerance in Celiac Disease. Clin Transl Gastroenterol 11, e00274, doi:10.14309/ctg.0000000000000274 (2020).

106 Capron, M. et al. Safety of P28GST, a Protein Derived from a Schistosome Helminth Parasite, in Patients with Crohn’s Disease: A Pilot Study (ACROHNEM). J Clin Med 9, doi:10.3390/jcm9010041 (2019).

